# Task-adapted biological foundation models uncover perturbation-centric representations

**DOI:** 10.64898/2026.06.30.735584

**Authors:** Elena Pareja-Lorente, Patrick Aloy

**Affiliations:** Institute for Research in Biomedicine (IRB Barcelona), The Barcelona Institute of Science and Technology, Barcelona, Catalonia, Spain; Institució Catalana de Recerca i Estudis Avançats (ICREA), Barcelona, Catalonia, Spain

**Keywords:** Foundation models, Perturbation biology, Chemical and genetic perturbations, Representation learning, MoA prediction, Target identification

## Abstract

Foundation models have emerged as powerful tools for learning transferable representations of biological systems, yet their latent spaces are typically optimized to capture cellular state rather than the effects of perturbations. Here, we demonstrate that a biological foundation model can be repurposed to learn a fundamentally different representation by changing its learning objective. We fine-tuned scGPT, a transformer pre-trained on over 30 million single-cell transcriptomes, on more than three million LINCS L1000 perturbation profiles using a supervised objective that predicts perturbation identity. This transformed the latent space into a perturbation-centric representation that aligned transcriptional responses induced by the same chemical or genetic perturbation across heterogeneous experimental conditions. Fine-tuned embeddings substantially outperformed both gene expression profiles and the original pre-trained model, recovering 85–100% of perturbations within the top 100 nearest neighbors and increasing perturbation classification accuracy from 10–19% to 25–49%. Remarkably, although the model was trained exclusively to recognize perturbation identity, the learned representation spontaneously captured orthogonal biological relationships never provided during training, including chemical similarity (AUROC up to 0.81), mechanisms of action (Hit@10 up to 100%), compound–target relationships (AUROC up to 0.74), and functional relationships between genetic perturbations. The resulting embedding space enabled mechanism-of-action annotation of nearly 12,000 previously uncharacterized compounds, prioritization of target-related chemical–genetic associations, and contextualization of unseen perturbations and external transcriptomic datasets. Together, our results establish objective-driven adaptation as a general strategy for repurposing biological foundation models to learn reusable representations of complex biological phenomena.

## Introduction

A defining feature of foundation models is that their latent representations are shaped not only by pretraining data but also by the objectives used during adaptation. Whether this principle can be exploited to learn entirely new biological representations remains largely unexplored.

The perturbation of a biological system through chemical compounds, genetic modifications, or environmental manipulations is often a powerful way to interrogate cellular functions^1^. Of all possible readouts, gene expression offers a high-dimensional description of cellular states following perturbation, capturing coordinated molecular responses^2^. Understanding how such perturbations reshape transcriptional states is central to uncovering molecular mechanisms, elucidating disease processes, and guiding therapeutic discovery^3,4^.

Gene expression-based perturbation profiling has been implemented at scale through large initiatives such as the Connectivity Map^5^ and the Library of Integrated Network-based Cellular Signatures (LINCS)^6^. More recent efforts include Chemical-Induced Gene expression Signatures (CIGS)^7^ and perturbational single-cell datasets like Tahoe^8^ and X-Atlas/Pisces^9^. While single-cell perturbation atlases hold considerable promise^10,11^, LINCS remains the most comprehensive and systematic perturbation resource available, as it contains over three million gene expression profiles generated across hundreds of cell lines, covering chemical and genetic perturbations induced at multiple doses and measures in different time points. By standardizing transcriptional measurements and representing perturbation effects as gene expression signatures, LINCS has enabled diverse applications, including mechanism of action discovery^12^, drug repurposing^13,14^, target identification^15^, and perturbation effect prediction^16–18^.

Despite its scale and potential, extracting generalizable biological signals from LINCS remains challenging. Multiple studies have shown that cell line-specific effects, batch variability, and platform-dependent biases often dominate transcriptional responses, obscuring the intrinsic effect of the perturbation itself^19–21^. As a result, gene expression signatures induced by the different perturbations in the same cellular context often look more similar than those induced by the same perturbation assayed in different cell lines. Recent cross-dataset analyses further showed that transcriptional responses induced by the same compound can exhibit weak agreement across experimental settings, in some cases falling below similarities observed between different compounds measured in the same context^22^. Together, these observations indicate that perturbation-specific signals are often masked by contextual and technical effects, limiting the robustness of perturbation representations across experimental conditions and constraining downstream analyses based directly on raw gene expression data^23–27^.

These challenges have motivated the development of representation-learning approaches that transform noisy transcriptional profiles into more informative latent spaces. Such representations can improve phenotype-based drug discovery^21^, perturbation effect prediction^28^, and target identification^15^ by encoding perturbation responses in a latent space that reduces technical biases and facilitates downstream predictive modeling. Together, these studies highlight the value of moving beyond direct comparison of gene expression or differential expression signatures toward more informative perturbational representations^22^.

Recently, foundation models pre-trained on large-scale datasets have demonstrated strong capacity for learning transferable representations in fields such as natural language processing^29^ and computer vision^30^. In biology, transformer-based foundation models, including Geneformer^31^, scGPT^32^, and scFoundation^33^, have been pre-trained on single-cell RNA-seq data coming from millions of cells to learn representations of cellular states. These models have been shown to capture gene-gene dependencies and co-expression patterns and produce embeddings that can be transferred across downstream tasks^32^. However, the pre-trained versions of these models are not able to consistently outperform simpler, domain-specific approaches in predictive settings^34–37^, suggesting that their utility may depend on task-specific fine-tuning and objective design.

Here, we show that a biological foundation model can be systematically repurposed by changing its learning objective, transforming a cell-state representation into a perturbation-state representation, which enables direct comparisons across conditions and their readily integration into predictive models^21,38,39^. We assess whether the new embedding space is able to separate perturbation-specific transcriptional effects from contextual variability. More specifically, we evaluate if the fine-tuned model is able to capture biologically meaningful relationships not explicitly encoded during training, such as mechanisms of action (MoA), genetic and chemical target perturbations, and the contextualization of new perturbation experiments within a unified perturbational space.

## Results and Discussion

### Learning perturbation representations from transcriptional responses

The main objective of this study was to learn perturbation-centric representations from transcriptional data, ensuring that embeddings corresponding to the same perturbation remain consistent across heterogeneous contexts and distinct from embeddings of unrelated perturbations. We started from scGPT^32^, a transformer-based foundation model pre-trained on over 30 million single-cell RNA-seq profiles from diverse tissues. scGPT employs a masked modeling objective, similar to natural language models^29,30,40^, to capture biological background and gene-gene dependencies, facilitating knowledge transfer to diverse downstream applications^41^. We then adapted this model to accept bulk perturbational data, fine-tuning scGPT on the LINCS L1000 Level 3 dataset^6^, which contains over three million gene expression profiles, measuring 978 landmark genes (Figure 1).

**Figure 1.**
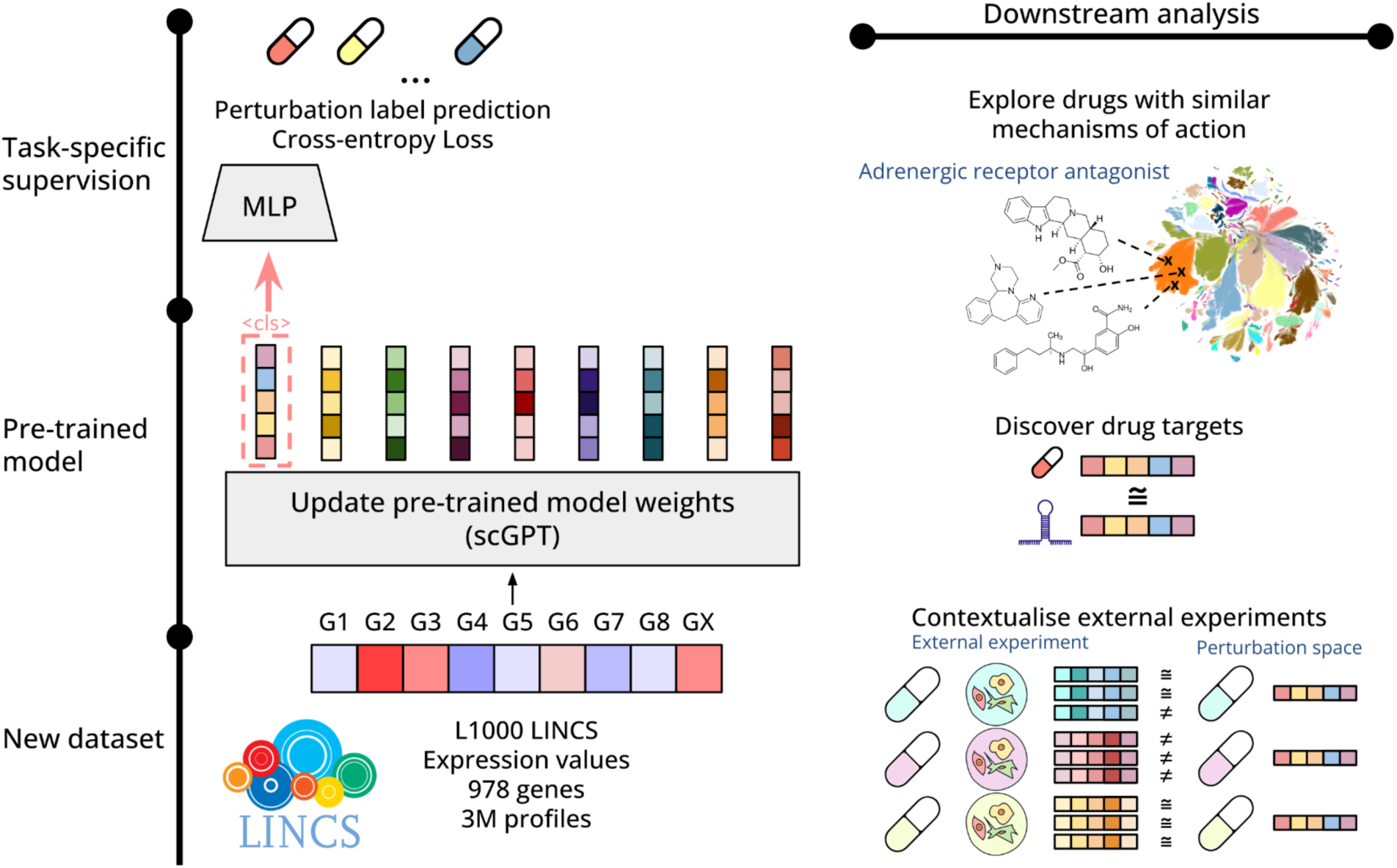
Schematic overview of the pipeline used in the Perturbation Representation study. We fine-tuned scGPT on the LINCS L1000 dataset to predict perturbation labels. The fine-tuning objective promotes alignment of embeddings from profiles corresponding to the same perturbation, thus capturing shared transcriptional signatures. The resulting embedding space enables multiple downstream applications, including (i) identifying compounds with shared Mechanisms-of-Action, (ii) prioritizing targets by comparing genetic and chemical perturbations, and (iii) contextualizing external datasets within a unified perturbational space.

We trained the model using a supervised perturbation-label classification objective that encouraged embeddings derived from profiles corresponding to the same perturbation to cluster in the latent space. We coupled the transformer encoder to a multilayer perceptron (MLP) classifier and optimized it using a cross-entropy loss to predict perturbation identity (see Methods for further details). By forcing the model to discriminate among a large number of perturbation labels, this objective promotes the separation of perturbation-specific transcriptional effects from contextual variability, including cell line identity, batch effects, and experimental conditions. Our training strategy produced a perturbation-centric embedding space that encodes transcriptional responses at perturbation level rather than at the experimental context level. These embeddings provided the basis for subsequent analyses, including mechanism of action inference, target-related signal, and contextualization of new perturbational datasets within a shared perturbational space.

#### Validation of the perturbational embedding space

To evaluate whether our fine-tuning strategy produced perturbation-centered representations able to overcome the cell context bias of the original model, we compared three embedding spaces: (i) pre-trained scGPT embeddings generated from LINCS profiles using the original scGPT model; (ii) embeddings from the model fine-tuned on the full LINCS dataset; and (iii) embeddings from the model fine-tuned on a subset of high-quality (HQ) LINCS profiles (see Methods).

We first visualized each representation using UMAP projections (Figure 2A, Supplementary Figure 1A-C). For both normalized (Level 3) gene expression (Gex) profiles and pre-trained scGPT embeddings, as expected, cell-line identity predominantly organized the representations, forming large, well-defined clusters. In contrast, embeddings derived from the fine-tuned models, both full and HQ, exhibited a reduced cell-dependent structure and instead formed a more continuous landscape with heterogeneous clusters. These results indicated that fine-tuning fundamentally shifted the latent space from encoding cellular context toward encoding perturbation identity. To quantify these observations, we computed pairwise cosine distances between profiles in each representation (Figure 2B, Supplementary Figure 1D). We compared gene expression profiles and embeddings from: (i) same cell line, different perturbations, (ii) same perturbation across different cell lines, and (iii) random perturbation-cell pairs. In Gex and pre-trained embeddings, distance distributions for the same perturbation and random pairs largely overlapped, indicating that perturbation identity marginally contributes to similarity between the profiles. In contrast, fine-tuned embeddings showed clear separation between these distributions: profiles corresponding to the same perturbation are significantly closer in latent space (Mann-Whitney U p-value = 0.0 for both full and HQ sets), while random pairs remained distant. We next assessed intra-perturbation label agreement to determine whether fine-tuning increased the coherence of profiles representing the same perturbation in comparison to original Gex profiles (Figure 2C). Using percentile-normalized within-perturbation distances, fine-tuned models exhibited lower average intra-perturbation distances than Gex, whereas the pre-trained model showed coherence comparable to Gex. The percentage of perturbation classes showing statistically significant intra-cluster coherence reached 26.96%, 77.24%, and 61.64% for the pre-trained, full fine-tuned, and HQ fine-tuned spaces, respectively. Analyses of additional metadata features, including cell line and plate, confirmed that fine-tuned embeddings systematically reduced confounding effects while preserving biologically meaningful structure (Supplementary Figure 2).

**Figure 2.**
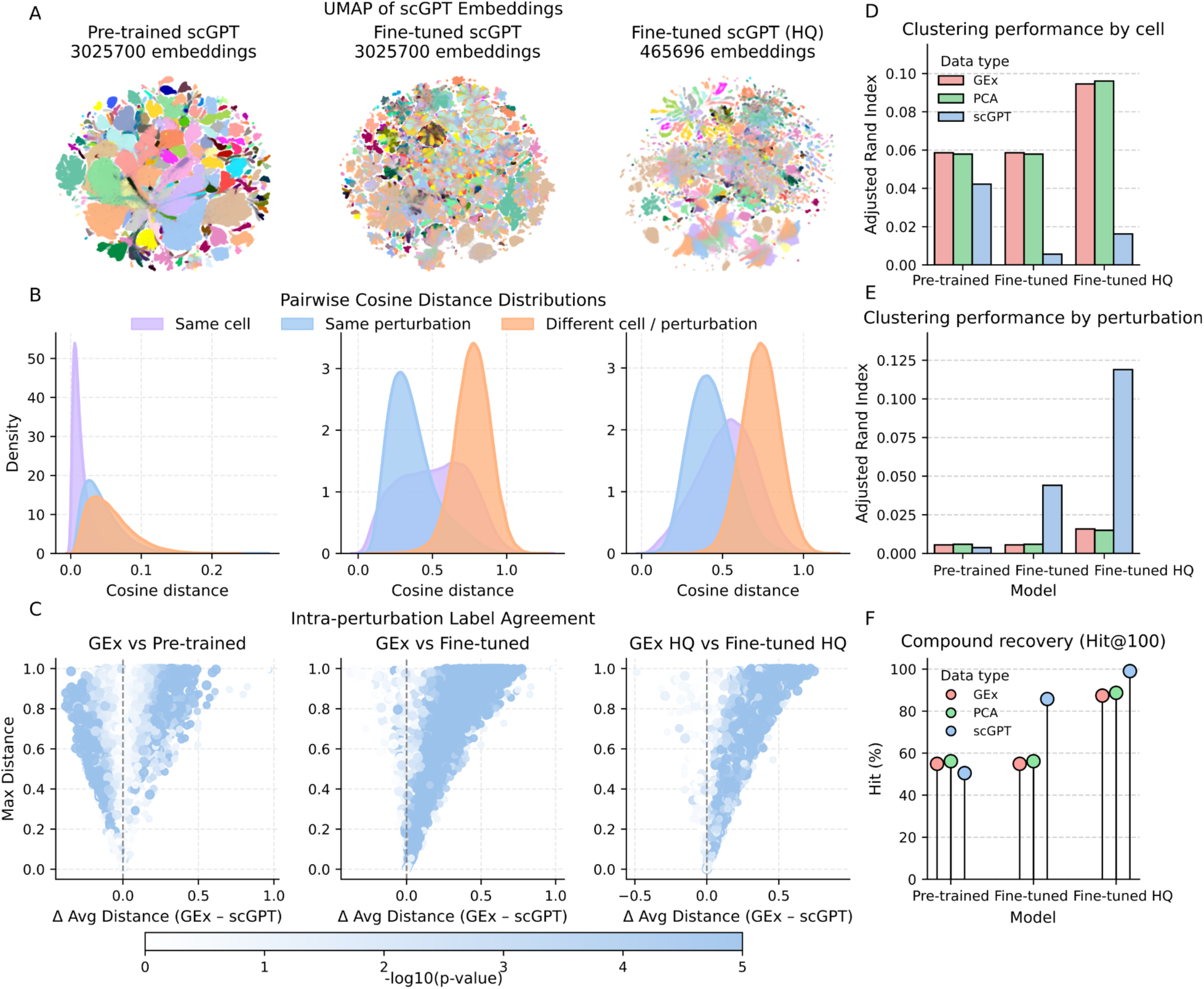
Perturbational space exploration. We constructed three perturbational spaces using a pre-trained scGPT model, a full fine-tuned version, and a fine-tuned model restricted to high-quality LINCS data. **A.** UMAP representation of scGPT embeddings derived from the LINCS L1000 (Level3) dataset. Colors represent the cell line of origin. **B.** Pairwise cosine distance distributions for scGPT embeddings grouped by same cell (purple), same perturbation (blue), or random (orange) pairs. **C.** Intra-coherence scatterplots. Each point represents a perturbation label; the x-axis shows the difference in average percentile-normalized within-perturbation cosine distance between Gex profiles and scGPT embeddings (left = tighter in Gex, right = tighter in scGPT embeddings), and the y-axis shows the maximal within-perturbation distance. Point size reflects cluster size, and color (-log₁₀ p) indicates the significance of the difference. **D-E.** Adjusted Rand Index (ARI) from clustering by cell line (D) or perturbation label ©. We compare Gex profiles (red), PCA representation (green) and scGPT embeddings (blue). **F.** Recovery of compounds sharing perturbation label quantified as Hit@100 (% of queries for which at least one of the top-100 nearest neighbors shares perturbation identity).

To further assess the biological relevance of our perturbation representations, we compared our embeddings against principal component analysis (PCA) representations of equal dimensionality (512 components). Using K-means clustering, we evaluated cluster agreement using the Adjusted Rand Index (ARI), Normalized Mutual Information (NMI), cluster dominance, cluster purity, and Silhouette score (Supplementary Table 1, Figure 2D-E). Full fine-tuned and HQ fine-tuned embeddings substantially outperformed both Gex and PCA in perturbation-specific clustering metrics (e.g., ARI and NMI), while achieving lower clustering scores for contextual factors such as cell line or time point. For example, ARI increased from 0.006 and 0.016 in Gex to 0.04 and 0.12 in the full and HQ fine-tuned embedding spaces, respectively. Similarly, cluster dominance for perturbation labels increased from 6% and 9% in Gex to 19% and 30% in the full and HQ fine-tuned embedding spaces. We further examined local neighborhood consistency by computing the proportion of profiles for which at least one of the *K* nearest neighbors (for *K* = 1, 5, 10, 100) shared the same perturbation label (Figure 2F, Supplementary Figure 3). Fine-tuned embeddings retrieved correct neighbors more often, with 85-100% of profiles recovering at least one true neighbor within the top 100, compared to <60-90% in the Gex space for the full and HQ datasets, respectively. Precision across *K* values consistently favored the fine-tuned models, confirming improved encoding of perturbation identity (Supplementary Figure 4).

Finally, to evaluate whether the learned embeddings facilitated perturbation classification, we trained supervised classifiers using either normalized gene expression (Gex) profiles or embedding representations as input features to assess how effectively each representation captured perturbation-specific information in a predictive setting. The task was intrinsically challenging because the full dataset contained 75,666 perturbation labels and the HQ dataset contained 32,252 labels. Nevertheless, classifiers trained on fine-tuned embeddings substantially outperformed classifiers trained directly on Gex profiles under identical train-test splits (Supplementary Figure 5A). Accuracy increased from 10.14% to 25.19% in the full dataset and from 18.97% to 48.74% in the HQ dataset. Performance varied across perturbation types, with gene silencing perturbations achieving the highest accuracy, consistent with their stronger and more reproducible transcriptional responses, as reflected by higher Transcriptional Activity Scores (TAS) (Supplementary Figure 5B).

#### Emergent biological organization in the perturbation latent space

A central question was whether optimizing the model exclusively for perturbation identity would produce a representation specialized only for this task, or whether broader biological organization would emerge. Having established that the fine-tuned embeddings captured perturbation-specific transcriptional effects, we next assessed whether the learned embedding space also encoded orthogonal biological relationships that we did not encode during training.

We first evaluated whether the fine-tuned embeddings recapitulated chemical similarity, given that structurally similar compounds often share biological effects. We formulated this exercise as a binary classification task, defining highly similar chemical pairs, those within the top 0.1 percentile of Tanimoto similarity, as positives and all remaining pairs as negatives. We computed cosine distances between embeddings and evaluated performance using ROC curves (Methods). Fine-tuned embeddings significantly outperformed Gex, achieving AUROCs of 0.81 and 0.78 for the full and HQ spaces, respectively, compared with 0.64 and 0.58 for Gex (Figure 3A).

**Figure 3.**
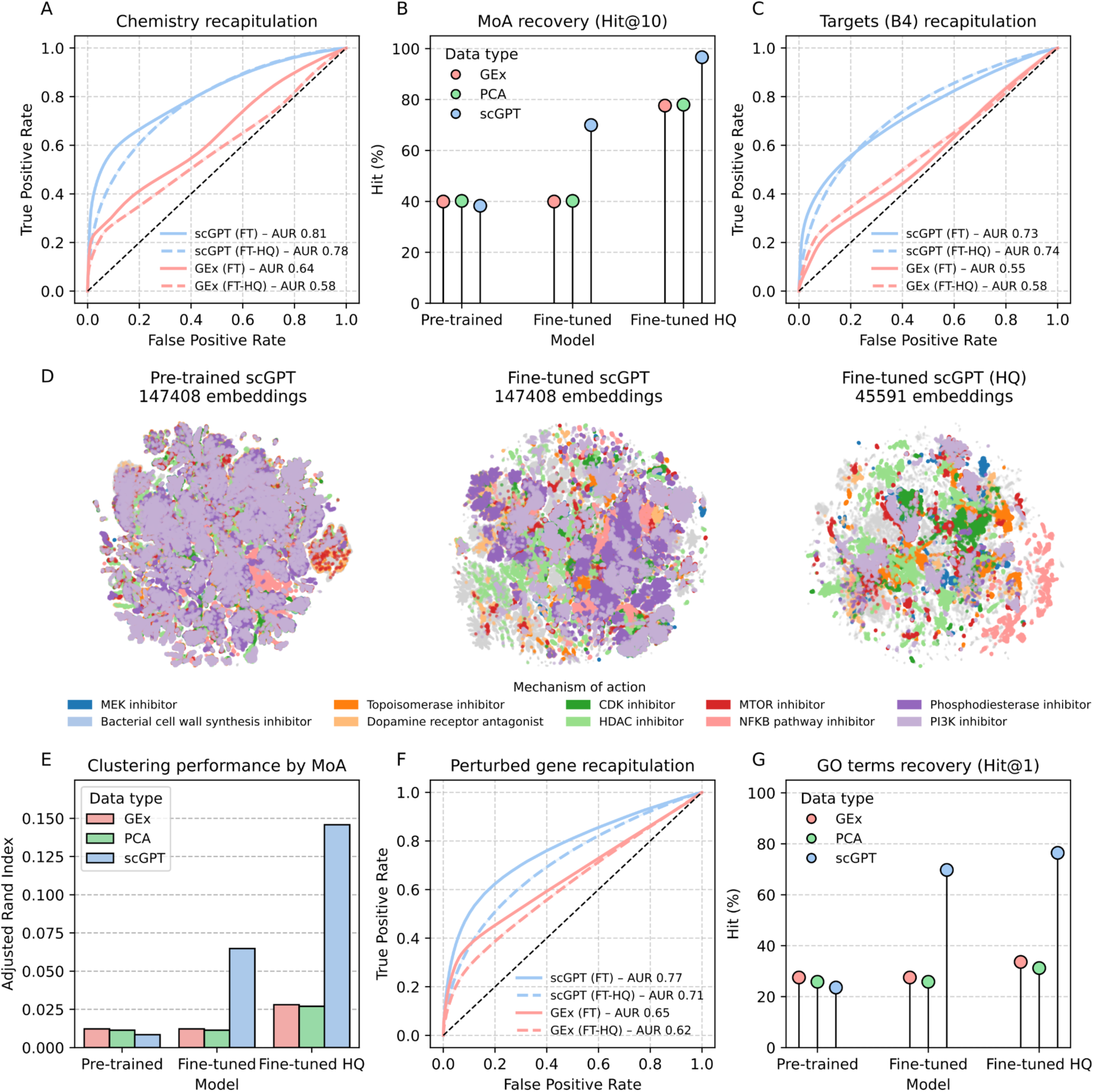
Recovery of orthogonal relationships by scGPT embeddings. **A.** Recapitulation of chemical similarity as a binary classification task. Positive pairs correspond to compound pairs within the top 0.1% of the Tanimoto similarity distribution, and we treated all remaining pairs as negatives. We used cosine distance between scGPT embeddings to quantify similarity. We show performance for gene expression profiles (Gex, red) and scGPT embeddings (blue) derived from the fine-tuned model (FT, solid lines) or from the fine-tuned model trained exclusively on high-quality data (FT-HQ, dashed lines). Legends report mean area under the ROC curve (AUROC). **B.** Recovery of compounds sharing Mechanism-of-Action (MoA) quantified as Hit@10 (% of queries for which at least one of the top-10 nearest neighbors shares MoA). We compare Gex profiles (red), PCA representation (green) and scGPT embeddings (blue). **C.** Recapitulation of protein targets. We show ROC curves for recovering compounds annotated to the same target using Gex or scGPT embeddings from FT and FT-HQ models, as in panel A. **D.** UMAP representation of scGPT embeddings colored by MoA classes with the largest number of annotated compounds. **E.** Clustering agreement by MoA, measured using the Adjusted Rand Index (ARI). **F.** ROC curves for recovery of the perturbed gene (from specific shRNA perturbations) using Gex or scGPT embeddings from FT and FT-HQ models, colored as in panel A. **G.** Recovery of Gene Ontology (GO) terms associated with perturbed genes (shRNA perturbations), quantified as Hit@1 (percentage of queries for which the top-1 nearest neighbor shares at least one GO term).

To further characterize the biological information encoded in the embeddings, we extended the Hit@k analysis (Figure 2F) to shared mechanism of action annotations. Fine-tuned embeddings substantially improved recovery relative to gene expression (Gex) profiles, increasing hit rates from approximately 40% to 70% in the full fine-tuned model and from approximately 80% to nearly 100% in the HQ fine-tuned model (Figure 3B). We then assessed whether the embeddings also reflected compound target information. Using the Chemical Checker (CC) “B4 targets” signatures^13,42^, which describe known protein targets for thousands of bioactive molecules (including 15,535 present in LINCS), we repeated the classification framework described above. Fine-tuned embeddings outperformed Gex (Figure 3C), indicating that the learned perturbation space captured relationships between compounds based on their target profiles.

We then explored whether these biological relationships were reflected in the global organization of the embedding space. We visualized the embeddings in two dimensions and colored profiles according to their MoA annotations (Figure 3D). Whereas gene expression profiles and pre-trained embeddings exhibited diffuse organization by MoA (Supplementary Figure 6A), the fine-tuned embeddings produced more coherent clusters corresponding to known pharmacological classes, indicating that the learned representations captured underlying biological mechanisms. Distance analyses comparing same-MoA pairs with random pairs revealed modest but significant shifts in the embedding spaces (Supplementary Figure 6B). This effect became more pronounced when evaluated through clustering-based metrics. The Adjusted Rand Index (ARI) for MoA increased from 0.012-0.028 in Gex to 0.065-0.15 in the fine-tuned embeddings (Figure 3E, Supplementary Table 1). In addition, intra-label coherence analyses showed that 79.93% (full) and 69.50% (HQ) of MoA classes exhibited tighter within-class similarity in the embedding space than in Gex (Supplementary Figure 7). Together, these results indicated that the fine-tuned embeddings encoded meaningful aspects of compound biological activity despite the absence of MoA or chemical information during training.

To contextualize these findings relative to previous representation-learning approaches for LINCS data, we evaluated our embeddings using the perturbagen class (PCL) prediction framework proposed in a previous VAE-based study^43^, where VAE-derived latent representations achieved performances comparable to gene expression (Gex). In contrast, logistic regression models trained on our scGPT embeddings substantially outperformed both Gex and PCA representations in the same evaluation setting. The full and HQ fine-tuned embeddings achieved accuracies of 0.591 and 0.849, respectively, compared with 0.393 and 0.732 for Gex (Supplementary Figure 8).

Beyond chemical perturbations, we also evaluated representation of genetic perturbations. During training, we treated each shRNA probe as an independent distinct perturbation so the model could not explicitly learn that multiple probes targeted the same gene. We therefore performed an ROC-based classification analysis in which we defined positive pairs as profiles derived from different shRNAs targeting the same gene. Fine-tuned embeddings significantly improved recovery compared with Gex (Figure 3F), consistent with prior observations that raw expression responses to gene knockdown are noisy and probe-dependent^6,44–47^. Finally, we examined whether embeddings captured functional relationships between the genes. Using Gene Ontology (GO) annotations associated with perturbed genes, we again extended the Hit@k analysis to shared GO term annotations. Fine-tuned embeddings achieved nearly 80% recovery, compared with less than 40% in gene expression (Gex) space (Figure 3G), indicating that the model captured functional gene-gene relationships.

Together, these results suggested that fine-tuned scGPT embeddings not only preserved perturbation identity but also organized the latent space according to biologically meaningful relationships between perturbations. This structure highlighted the ability of foundation models to infer biological associations across perturbation profiles, providing a robust representation for downstream analysis.

### Unlocking biological insights from the perturbation space

#### Nearest Neighbor-based Imputation of Mechanisms-of-Action

Given that the embedding space organizes compounds according to their mechanisms of action, we assessed whether this structure could be used to infer MoA annotations for unannotated compounds in LINCS. Thus, for each unannotated compound in the full fine-tuned embedding space, we retrieved its top 50 annotated nearest neighbors based on cosine distances. We converted distances into similarity scores using a softmax function (i.e. weighting the proximity to the annotated compounds) and aggregated these scores at the MoA level by summing the contributions of neighbors sharing the same annotation (see Methods). For each unannotated compound, we averaged MoA probabilities across all its profiles and assigned the MoA with the highest aggregated probability as the predicted label. We used this aggregated probability as a confidence score. Overall, we could initially assign a MoA to 31,240 previously unannotated compounds, spanning 280 distinct mechanisms. The resulting MoA distribution showed moderate agreement with the distribution observed in annotated compounds (Spearman correlation = 0.51), suggesting that the imputation partially preserves the global structure of MoA classes (Supplementary Figure 9A).

To assess accuracy of the predictions, we examined the distribution of confidence scores and prioritized high-confidence predictions (Figure 4A). We observed that confidence scores varied across MoA classes, with HDAC inhibitors among the highest-confidence predictions (Supplementary Figure 9B). This observation is consistent with HDAC inhibitors tending to induce stronger transcriptional responses than many other MoA classes, as reflected by their higher TAS distributions (Supplementary Figure 9C). We manually inspected top-confidence predictions and found multiple biologically plausible annotations supported by literature. For example, we identified known HDAC inhibitors such as oxamflatin^48^, HC-toxin^49^, and fluoro-SAHA^50^; protein synthesis inhibitors such as cephaeline and emetine-HCL^51^; ATPase inhibitors such as sarmentogenin, digitoxigenin^52^, and peruvoside^53^; and HSP inhibitors such as XL-888^54^ (Figure 4B).

**Figure 4.**
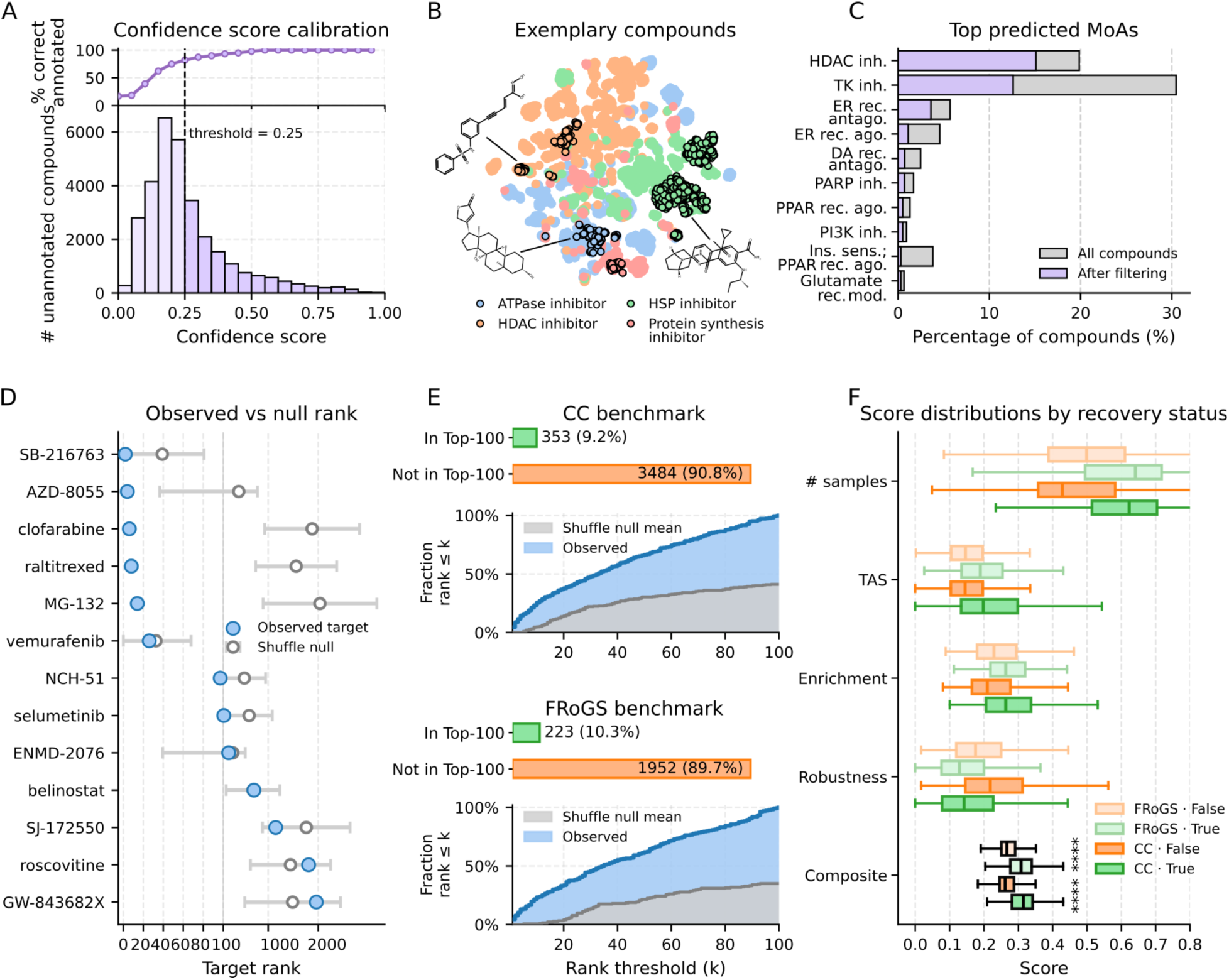
Exploring the fine-tuned perturbational space to uncover novel relationships. **A.** Calibration of confidence scores for MoA prediction. Upper panel: cumulative accuracy of MoA predictions across increasing confidence thresholds in the leave-one-compound-out benchmark. Lower panel: distribution of confidence scores across unannotated compounds. The dashed line indicates the selected confidence threshold (0.25). **B.** t-SNE representation of compound embeddings showing selected examples of MoA imputation. Annotated compounds are shown as filled points, whereas previously unannotated compounds are outlined in black and colored according to their predicted MoA. Chemical structures shown from upper left to lower right: oxamflatin, digitoxigenin, and XL-888. **C.** Distribution of the most frequent predicted MoAs before and after confidence filtering. **D.** Benchmark evaluation of compound-target recovery. For each benchmark compound, the rank of the annotated target based on compound-shRNA embedding distances is compared against a shuffled-label null distribution. Lower ranks indicate better recovery. **E.** Recovery of known targets among the top 100 predicted genes across benchmark datasets (Chemical Checker and FRoGS). Upper panel: number of compounds with at least one recovered target. Lower panel: cumulative recovery as a function of best target rank compared with the shuffled null expectation. **F.** Distribution of confidence features stratified by target recovery status across the Chemical Checker and FRoGS benchmarks. Boxplots show the number of QC-passing profiles, transcriptional activity score (TAS), enrichment score, robustness score, and final composite confidence score.

To quantitatively evaluate the accuracy of the MoA imputations and to define a confidence threshold, we benchmarked the approach on annotated compounds. This is, for each compound belonging to MoA classes with multiple members, we iteratively removed all its profiles from the dataset and predicted its MoA using the same nearest-neighbor procedure (see Methods). We observed a strong positive relationship between prediction confidence and accuracy, indicating that the confidence score provides a useful ranking of prediction reliability, with higher-confidence predictions showing higher proportions of correct assignments in the leave-one-compound-out benchmark (Figure 4A). Based on this analysis, we selected a confidence threshold of 0.25, corresponding to 81.92% correct predictions. Applying this threshold to the unannotated set, we retained 11,787 compounds with high-confidence MoA annotations, spanning 69 distinct MoA classes. The most frequently assigned MoAs included HDAC inhibitors and Tyrosine Kinase inhibitors, although their relative overrepresentation decreased after filtering (Figure 4C). All the high-quality inferred annotations, including the assigned MoA, confidence score, agreement between compound samples, and average TAS for each unannotated compound are available in Supplementary Table 2.

#### Linking chemical and genetic perturbations through target and mechanistic similarity

A key challenge in perturbation biology is linking chemical and genetic perturbations acting on the same target. Despite the intuitive expectation that pharmacological inhibition and genetic perturbation of the same gene should induce similar transcriptional responses, multiple previous studies have reported weak or inconsistent correlations between these perturbation types. This reflects the biological complexity of compound polypharmacology, compensatory mechanisms, and context-specific responses^1,15,55,56^. We therefore evaluated whether the learned perturbation embedding space improves the alignment between chemical and genetic perturbations targeting the same gene.

We first analyzed a benchmark set of 13 compounds with well-curated target annotations in LINCS that previous studies reported to exhibit relatively conserved transcriptional responses across perturbation modalities^6^. For each compound, we computed cosine distances between compound embeddings and all shRNA embeddings. Then, for every compound-profile embedding, we retrieved the top 100 nearest shRNA embeddings and aggregated the results at the compound level. Specifically, for each gene in the selected shRNA neighborhood, we computed a robustness score reflecting how frequently shRNAs targeting that gene appeared among the top 100 shRNA neighbors across all profiles of the compound. We then ranked genes according to this robustness score to generate a compound-level target ranking.

Using this strategy, we recovered the annotated target among the prioritized genes for 7 out of the 13 benchmark compounds. However, when we compared these results against random expectation, we found that recovery for 2 of these compounds was not substantially better than expected by chance (Figure 4D). We manually inspected the ranked gene lists, and we found that this benchmark likely underestimates biologically meaningful matches, as several compounds recovered alternative known targets or functionally related genes rather than the specific annotated target used for evaluation. For instance, we could not recapitulate the annotated target genes for ENMD-2076 (AURKB) and roscovitine (CDK9), but we found alternative known related targets among the highest-ranked genes (ENMD-2076: AURKA rank #16; roscovitine: CDK2 rank #43). Additionally, for another two compounds, namely selumetinib (MAP2K1) and GW-843682X (PLK1), highly ranked genes were functionally related to the annotated target through shared pathways: selumetinib predictions were enriched in MAPK cascade-related terms, whereas GW-843682X predictions were associated with cell proliferation and apoptotic processes. These observations suggest that strict one-compound/one-target evaluation schemes may overlook biologically relevant relationships in perturbational data.

To broaden our observation to a systematic evaluation, we expanded the analysis using compound-target annotations from the Chemical Checker^13^. After mapping compounds and targets to the LINCS perturbational space, we assigned 3,837 compounds onto 1,331 targets. For each compound, we evaluated whether at least one known active target appeared among the top 100 highest-ranked genes. Using this criterion, we successfully recovered targets for 353 compounds, including 9 out of the 12 mapped compounds from the previous benchmark set. Importantly, the cumulative rank distributions showed that correctly recovered targets were strongly enriched among top-ranked genes relative to the shuffled null expectation, demonstrating that successful predictions were not only recoverable within the top 100 candidates but were systematically prioritized near the top of the ranked lists (Figure 4E). We observed similar behavior in an independent benchmark provided by FRoGS^15^, containing 2,175 compounds, where we recovered targets for 223 of them (Figure 4E). Together, these results confirm that linking chemical and genetic perturbations remains a challenging problem and that correspondence between perturbation modalities is far from trivial. Nevertheless, the embedding space captured partial but biologically meaningful relationships between compounds and genetic perturbations, including pathway-level associations and target-related signals beyond direct target matching. However, the variability in recovery performance across compounds also indicated that not all associations reflected robust mechanistic relationships, motivating the development of a confidence scoring framework to distinguish meaningful biological signal from spurious correlations.

To better understand why some compound-target relationships were more consistently recovered than others, we examined whether recovery performance was associated with properties of the underlying compound profiles. We first considered the Transcriptional Activity Score (TAS), a LINCS metric reflecting the strength and reproducibility of perturbational signatures^25^. We observed that low-TAS embeddings tent to cluster together even when they represent unrelated perturbations, suggesting unspecific signatures most likely due to a lack of strong specific response (Supplementary Figure 10A). Indeed, the average cosine distance from low-TAS embeddings to random unrelated signatures is significantly lower than those computed from high-TAS embeddings (Supplementary Figure 10B). Moreover, in both benchmark datasets, compounds whose annotated targets were recovered at higher ranks tended to have higher TAS values, as reflected by a negative correlation between target rank and median compound TAS (Spearman ρ = -0.275 and -0.282 for FRoGS and Chemical Checker, respectively). Consistently, compounds for which the known target was recovered within the top 100 ranked genes showed higher TAS distributions than non-recovered compounds in both benchmarks (mean 0.19 vs 0.14 for FRoGS; mean 0.21 vs 0.14 for Chemical Checker; Figure 4F). We observed a similar, although weaker, association with the number of available compound profiles, with higher profile support associated with better target prioritization (Spearman ρ = -0.18 and -0.16 for FRoGS and Chemical Checker, respectively). Recovered compounds also showed higher profile-support distributions than non-recovered compounds in both benchmarks (mean 679 vs 224 for FRoGS; mean 587 vs 165 for Chemical Checker; Figure 4F). Together with the observation that some high-ranking gene lists captured pathway-level or mechanistic relationships even when the exact annotated target was not recovered, these results suggested that prediction reliability depends on multiple complementary factors, including transcriptional signal strength, profile support, and functional coherence. We therefore explored whether these features could be combined to identify higher-confidence compound-gene predictions.

To exploit these observations and prioritize higher-confidence predictions, we implemented a confidence scoring framework based on four complementary features: (i) the number of quality-control passing profiles per compound, capturing the stability of the aggregated prediction; (ii) the mean transcriptional activity score (TAS) of compound signatures, reflecting transcriptional strength and reproducibility; (iii) functional coherence among the top 100 predicted genes, quantified as the strongest enrichment across KEGG, Reactome, and Gene Ontology Biological Process annotations; and (iv) the mean robustness score of the top-ranked genes. By combining these features, we could significantly discriminate between target genes with close chemical and genetic perturbation embeddings from those that could not be related in the benchmark datasets (Figure 4F), indicating that we can estimate prediction confidence from sample quality, reproducibility, and functional consistency. Indeed, we observed that with a confidence threshold set to 0.35, over 1/3 of target inferences corresponded to correct gene recoveries (Supplementary Figure 9D-E), demonstrating that this score enriches for higher-confidence compound-target associations. We then applied this confidence threshold to all compounds in the embedding space to generate prioritized compound-gene association lists, retaining 1,510 compounds for which we predicted their potential targets. Supplementary Table 3 provides a prioritized lists of target genes for each compound, together with all associated confidence metrics.

#### Contextualization of unseen compounds and cellular contexts

To evaluate the ability of the model to contextualize within the perturbation embedding space gene expression profiles not observed during training, we designed two out-of-distribution settings using the HQ fine-tuned model. In the first setting, we excluded all gene expression profiles corresponding to 50 compounds from training, which were structurally diverse and representative of the chemical heterogeneity observed in the training set (Supplementary Figure 11). In the second setting, we excluded all profiles derived from a single cell line (A549), thereby assessing generalization across cellular contexts.

This set of 50 held-out compounds comprised 2,190 gene expression profiles. For these profiles, we generated embeddings using the new trained model and computed cosine distances to all embeddings from the training set using FAISS^57^. We evaluated contextualization accuracy by assessing whether the nearest neighbors corresponded to compounds sharing the same MoA. Using the predicted embeddings, more than 36% of the profiles recovered a nearest neighbor with the same MoA, compared with a random expectation of 1.56% (estimated using a hypergeometric model; see Methods). When expanding the neighborhood to the top 10 nearest neighbors, we recovered at least one compound with the same MoA for 53% of the profiles, compared with a random expectation of 13.83% (Figure 5A). We additionally evaluated contextualization using a clustering-based approach. We performed K-means clustering on the training embeddings, and we assigned the held-out compound embeddings to the predicted cluster. We assigned 50.4% of the predicted embeddings to clusters enriched for the correct MoA (Supplementary Figure 12A). These results indicate that the learned embedding space can place novel compounds into biologically meaningful neighborhoods even when their perturbation labels are absent from the training set.

**Figure 5.**
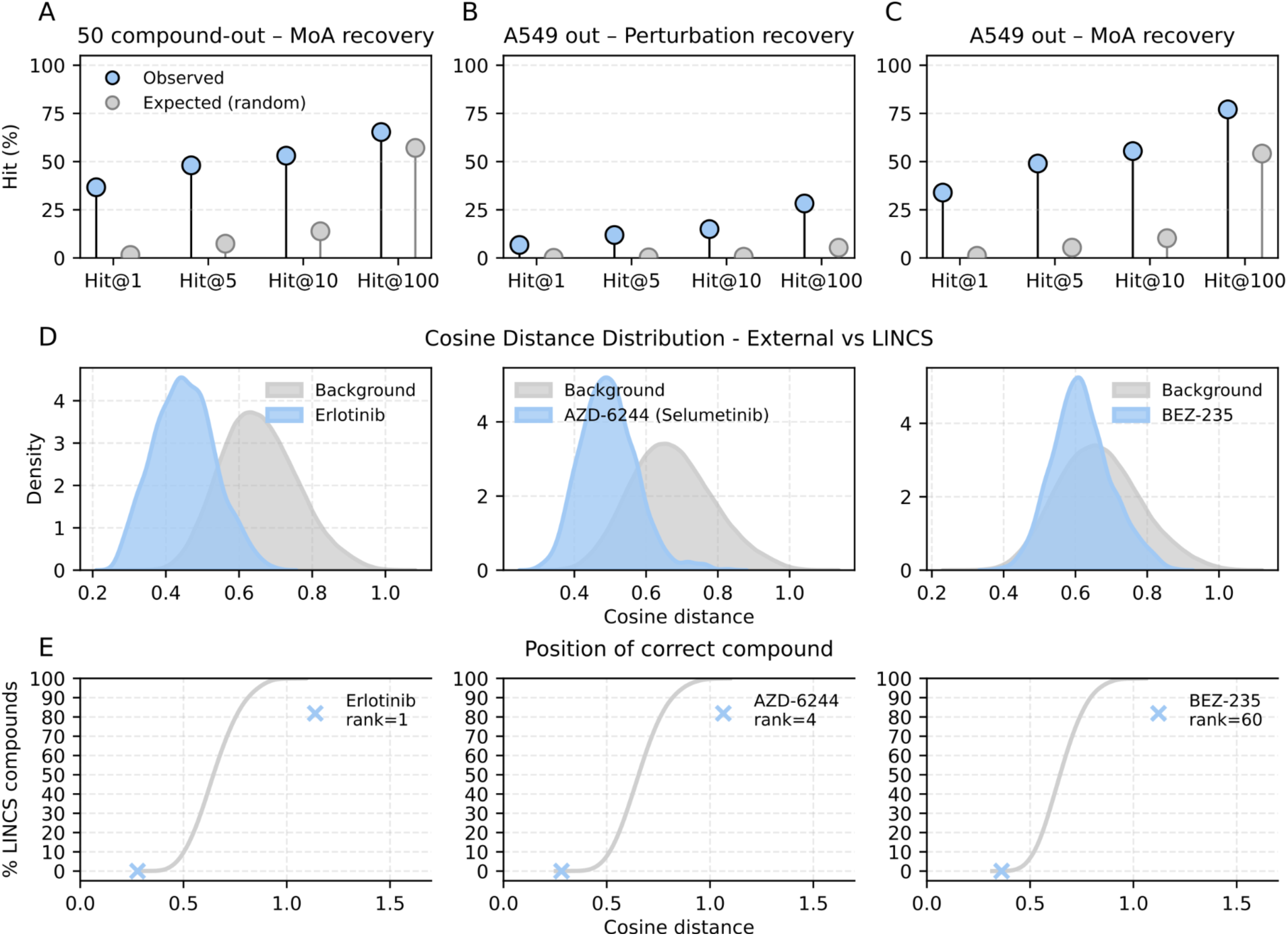
Generalization of HQ fine-tuned scGPT embeddings across contexts and external datasets. **A-C.** Contextualization performance of HQ fine-tuned scGPT embeddings under two hold-out settings, quantified as Hit@k (k = 1, 5, 10, 100). Results are shown for compound hold-out evaluated by MoA (**A**), and A549 cell-line hold-out perturbations evaluated at the level of exact perturbation labels (**B**) and by shared MoA (**C**). For each k, observed performance (blue) is shown alongside the expected performance under random retrieval (grey). **D.** Cosine distance distributions between embeddings derived from an external dataset (GSE51212) and LINCS compound embeddings for three representative compounds (Erlotinib, AZD-6244, and BEZ-235). For each compound, the background distribution corresponds to distances to all LINCS compounds, while the blue distribution corresponds to distances to the matching compound measured in LINCS. **E.** Empirical cumulative distribution of cosine distances from external perturbation embeddings to all LINCS compounds for the same three examples. The rank at which the first correctly contextualized LINCS compound is recovered is highlighted (cross).

We next evaluated contextualization across unseen cellular contexts using the A549 cell line, which was entirely excluded from training. In this setting, recovery of the exact perturbation label remained challenging: only 15% of A549 profiles recovered a nearest neighbor with the same perturbation label within the top 10 neighbors, compared with a random expectation of 0.65% (Figure 5B). However, when we assessed contextualization at the level of broader annotations, the performance of the embeddings improved substantially. Specifically, 34% of A549 compound-perturbed profiles recovered a nearest neighbor sharing the same MoA, and more than 55% recovered at least one MoA-consistent neighbor within the top 10, compared with random expectations of 1.1% and 10.24%, respectively (Figure 5C). Using the same clustering-based approach applied to held-out compounds, we found that 56.5% of the A549 embeddings were assigned to clusters enriched for the correct MoA (Supplementary Figure 12A).

To further improve contextualization across unseen cellular contexts, we explored two adaptation strategies. First, we fine-tuned a new high-quality model excluding all A549 perturbation profiles but retaining A549 DMSO controls, allowing the model to observe the cell line under baseline conditions. Second, we performed additional fine-tuning epochs using a reconstruction objective on A549 DMSO profiles. Neither strategy yielded substantial improvements over the non-adapted model (Supplementary Figure 12B), likely due to the limited number of available A549 control profiles relative to the full training set and the mismatch between reconstruction-based and perturbation-centric objectives. To better understand this limitation, we evaluated how well different cell lines were integrated into the embedding space as a function of the number of available profiles (Supplementary Figure 12C). As expected, cell lines represented by large numbers of profiles likely drive the structure of the perturbation embedding space, showing substantially better integration reflected by lower differences between within-cell-line and between-cell-line distances. In contrast, cell lines represented by fewer profiles (i.e. < 2,000) exhibited a clear tendency to remain dominated by cell line-specific effects. These observations support the idea that contextualization across unseen cellular environments remains particularly challenging when the model has not previously observed sufficient transcriptional variability associated with a given cellular background.

Together, these results support that the fine-tuned perturbation embeddings can place previously unseen perturbations into biologically meaningful neighborhoods, even when the corresponding compounds or cellular contexts were absent from training. Additionally, they also highlight the intrinsic difficulty of contextualizing perturbations across new cellular environments^58,59^.

#### Contextualization of external datasets

Finally, to evaluate whether the learned embedding space could contextualize gene expression data generated in a completely different setting and experimental pipeline (i.e. outside of LINCS), we performed a projection analysis using an independent dataset. We selected the GEO dataset GSE51212, which profiled transcriptional responses to three compounds, erlotinib, AZD6244, and BEZ235, in HCC827 human lung cancer cells^60^. All three compounds were present in LINCS, enabling direct comparison. Measurements were performed at 6 hours for all compounds, with additional time points (3, 12, and 24 hours) available for erlotinib.

We generated embeddings for the external gene expression profiles using the high-quality fine-tuned model and computed cosine distances to all embeddings from the high-quality LINCS reference set. We first compared distance distributions between embeddings corresponding to the same compound measured in LINCS and those corresponding to random compounds. This analysis revealed clear separation between the two distributions, indicating that embeddings derived from the external dataset were positioned closer to their corresponding perturbations in LINCS than expected by chance (Figure 5D). However, when we determined the rank at which the first correctly matched LINCS embedding appeared for each compound, we recovered the nearest matching embeddings at ranks 1, 4, and 60 for erlotinib, AZD6244, and BEZ235, respectively (Figure 5E). Thus, despite differences in experimental platform, cellular context, and time points, the model accurately projected external perturbation profiles into the learned embedding space and recovered their corresponding perturbations.

## Concluding remarks

In this study, we leveraged single-cell foundation models and perturbation-induced transcriptional responses to demonstrate that task-specific fine-tuning can substantially improve the representation of large-scale perturbation transcriptomic data. Rather than developing a predictor for a single downstream task, we sought to learn a reusable perturbation representation that could support diverse inference problems. Our results demonstrate that a task-adapted foundation model can serve as a general perturbation encoder from which multiple biological properties emerge. Indeed, perhaps the most striking observation is that no chemical descriptors, target annotations, mechanisms of action or pathway information were ever provided during training. And yet, these properties emerged naturally from a model trained only to discriminate perturbation identities.

At the same time, our results highlight important limitations of perturbation representation learning. Although LINCS represents a uniquely valuable resource, its profiles vary substantially in transcriptional strength, reproducibility, and experimental coverage. Weak or low-quality samples can cluster together in embedding space for reasons unrelated to shared biology, potentially generating spurious downstream associations. For this reason, embedding proximity should not be interpreted in isolation but together with confidence measures incorporating sample quality, transcriptional activity, replicate support, and functional coherence. Cellular context also remains a major source of variability. Although fine-tuning reduced cell line-driven structure, it did not eliminate it completely. Sparsely represented or biologically distinct cell lines remained more difficult to contextualize, suggesting that robust cross-context generalization requires broad representation of cellular backgrounds during training.

Future work should explore training strategies that more explicitly disentangle perturbation effects from contextual and technical covariates while incorporating additional modalities^61^ to extend these approaches toward perturbation effect prediction^62,63^. Contrastive learning, metric learning, adversarial objectives, or multi-task training could further improve representation robustness. More stringent out-of-distribution evaluations will also be important to define the generalizability of these representations. In addition, developing improved approaches to project single-cell perturbation atlases^64^ into LINCS-derived embedding spaces could provide a multi-resolution framework connecting bulk perturbation signatures with cell type-specific responses.

By releasing the fine-tuned embeddings, model, projection workflow, and high-confidence mechanism of action and compound-target prioritization predictions, we provide a reusable resource for the perturbation biology and drug discovery communities. This framework enables querying LINCS perturbations, contextualizing new experiments, identifying mechanistically related compounds, and prioritizing candidate targets or pathways for further validation. Because these embeddings provide compact and machine-learning-friendly representations of transcriptional responses while retaining biologically meaningful information, they may also support a broader range of downstream predictive tasks beyond those explored here. Our results establish task-adapted foundation models as a practical framework for transforming large-scale perturbational transcriptomics into interpretable, reusable, and hypothesis-generating maps of cellular response. More broadly, this work demonstrates that the utility of biological foundation models is determined as much by the downstream learning objective as by pretraining itself.

## Methods

### Datasets

We used the LINCS L1000 Level 3 dataset^6^ (2020 resource), which contains more than three million gene expression (Gex) profiles representing transcriptional responses to chemical and genetic perturbations. Each profile measures the expression of 978 landmark genes and derives from experiments performed across multiple cell lines, perturbation types, doses, and time points.

For the pre-trained and full fine-tuned models, we used all available Level 3 profiles downloaded from the CLUE data library (https://clue.io/data). Specifically, we downloaded the following matrices:

- *Level3_beta_trt_sh_n453175×12328.gctx,*
- *Level3_beta_trt_cp_n1805898×12328.gctx,*
- *Level3_beta_trt_oe_n131668×12328.gctx,*
- *Level3_beta_trt_xpr_n420583×12328.gctx,*
- *Level3_beta_ctl_n188708×12328.gctx,*
- *Level3_beta_trt_misc_n26428×12328.gctx*

These files collectively comprised 3,025,700 gene expression profiles across all perturbation types. We obtained compound metadata, including mechanism of action (MoA) and target annotations, from *compoundinfo_beta.txt*. We retrieved profile- and signature-level quality metrics from *instinfo_beta.txt* and *siginfo_beta.txt*.

To construct the high-quality (HQ) dataset used for HQ fine-tuning, we applied filtering criteria consistent with recommendations from the original LINCS publication. From *siginfo_beta.txt,* we retained only signatures satisfying the following criteria: *qc_pass == 1, is_hiq == 1, cc_q75 > 0.2, TAS ≥ 0.2, batch_effect_tstat_pct < 95.* After selecting signatures that met these criteria, we further filtered corresponding profiles using *instinfo_beta.txt,* retaining only profiles with: *qc_pass == 1.* This procedure resulted in a final HQ dataset comprising 465,696 gene expression profiles.

### Model architecture and fine-tuning procedure

We used scGPT^32^, a transformer-based foundation model pre-trained on 33 million single-cell RNA-seq profiles. scGPT employs a masked modeling objective adapted for gene expression data and produces contextualized embeddings for genes and cells. We downloaded the whole-human pre-trained checkpoint (“scGPT_human”) from the official repository (https://github.com/bowang-lab/scGPT/).

To adapt the pre-trained model to perturbational transcriptomic data, we fine-tuned scGPT using a multi-class perturbation classification objective. We treated each LINCS expression profile as an independent sample and assigned a perturbation label corresponding to its perturbagen identifier (e.g., compound ID, shRNA ID, overexpression construct). Profiles corresponding to different doses, replicates, or time points of the same perturbation shared the same label.

We mapped LINCS landmark genes to the gene vocabulary used by scGPT, successfully matching 968 out of 978 genes. We excluded genes without a corresponding token. We constructed model inputs following the preprocessing strategy implemented in the original scGPT framework. Briefly, we used quantile-normalized Level 3 expression values directly without additional normalization or log-transformation and discretized expression values into 51 bins, consistent with the original scGPT tokenization strategy.

We followed the fine-tuning procedure implemented in the official scGPT repository. First, we randomly split the dataset into training (90%) and validation (10%) subsets at the profile level. Because our objective was to learn a perturbation-centered embedding space rather than evaluate strict out-of-distribution predictive performance, we did not enforce perturbation-level separation between training and validation sets. This strategy allowed the model to observe heterogeneous contexts, doses, and replicates of the same perturbation during training and facilitated learning representations that capture shared perturbational effects across conditions.

We appended a three-layer MLP classifier to the output of the transformer encoder and optimized the model using a cross-entropy loss to predict perturbation identity (75,666 distinct perturbation labels in the full dataset and 32,252 in the HQ dataset). During fine-tuning, we updated all transformer parameters. The fine-tuning architecture used a 12-layer transformer encoder with hidden dimension 512, 8 attention heads, and dropout probability of 0.2. We trained the models using the Adam optimizer with a learning rate of 1 × 10⁻⁴ and batch size of 32. We disabled additional scGPT objectives, including masked value prediction and elastic cell similarity regularization, to focus optimization on perturbation classification. For the full LINCS dataset, we trained the model for 20 epochs, whereas for the HQ dataset we trained for 50 epochs. Training used automatic mixed precision and the fast transformer implementation provided in the original scGPT framework.

#### Embedding extraction

After training, we extracted embeddings for each gene expression profile using the *scg.tasks.embed_data()* function from the *scgpt package*. This procedure generated one fixed-length embedding vector of 512 dim per profile, corresponding to the final encoder representation. We constructed three embedding spaces: (1) Pre-trained embeddings obtained by applying the original scGPT_human checkpoint without fine-tuning to LINCS data, (2) full fine-tuned embeddings obtained after training on the complete LINCS dataset, and (3) HQ fine-tuned embeddings obtained after training on the filtered high-quality subset.

### Evaluation of perturbation embedding spaces

#### Dimensionality reduction and visualization

We projected embeddings and original expression profiles into two dimensions using Uniform Manifold Approximation and Projection (UMAP). For each representation (Gex, pre-trained embeddings, fine-tuned embeddings, and HQ fine-tuned embeddings), we first computed a k-nearest neighbor (kNN) graph using Scanpy (v1.10.3) with *sc.pp.neighbors* and default parameters. We then applied UMAP using *sc.tl.umap* to obtain two-dimensional projections for visualization. We colored UMAP projections using metadata annotations, including cell line identity, perturbation label, and MoA. For MoA visualization, we restricted the analysis to compounds with curated MoA annotations.

#### Pairwise distance computation

We computed pairwise cosine distances between selected samples in both the scGPT embedding spaces and original gene expression spaces. We implemented a controlled sampling framework to generate pairs of samples sharing specific features while differing in others. Specifically, we constructed three categories of sample pairs: (1) same perturbation, different cellular context: pairs of samples corresponding to the same perturbation identifier (*pert_id*) applied in different cell lines (*cell_iname*), (2) same cellular context, different perturbations: pairs of samples corresponding to the same cell line but different perturbations, and (3) randomly sampled pairs in which both perturbation identity and cell line differed.

For each perturbation and each cell line represented in the dataset, we randomly sampled up to 1,000 pairs per category. For the random category, we sampled 100,000 pairs to obtain a reference distribution. For each pair, we computed cosine distance between their vector representations. We compared the corresponding distance distributions using a two-sided Mann-Whitney U test.

#### Intra-Perturbation coherence

We evaluated intra-perturbation coherence, defined as the compactness of samples sharing the same perturbation label. To ensure balance evaluation across perturbations with heterogeneous sample sizes, we randomly selected 5,000 perturbations and sampled up to 10 profiles per perturbation (when available), using a fixed random seed. We restricted all analyses to this sampled subset.

For each perturbation, we computed all pairwise cosine distances between profiles belonging to the same perturbation in both Gex and embedding spaces. To account for differences in global distance scales between representations, we normalized within-perturbation distances using percentile ranks relative to the global pairwise distance distribution. Lower percentile values indicate tighter clustering. For each perturbation *k*, we computed the difference in average percentile-scaled within-group distance between Gex and embedding space (Δ_k_), where positive values indicate greater compactness in the embedding space. We compared within-perturbation distance distributions between representations using a two-sided Mann-Whitney U test and adjusted p-values using Benjamini-Hochberg FDR correction.

For visualization, we represented the difference in average within-group percentile-scaled distance (x_axis), the robust maximal dispersion (95th percentile of within-group distance; y-axis), point size proportional to the number of samples per perturbation and point color corresponding to -log10(FDR).

We repeated this analysis for additional metadata categories, including cell line, time point, concentration, RNA plate, and MoA for compound perturbations.

#### Clustering and evaluation metrics

For each model variant (pre-trained, full fine-tuned and HQ fine-tuned), we generated cluster labels in three feature spaces: (i) gene expression profiles (978 genes), (ii) PCA-reduced representations with dimensionality matched to scGPT embeddings (512 components), and (iii) scGPT embeddings.

Due to dataset size, we applied MiniBatchKMeans clustering (*sklearn*) with a batch size of 10,000 and a fixed random seed. We defined the number of clusters using the heuristic *K* = √*N*/2, where *N* denotes the number of samples in the corresponding dataset.

We quantified alignment between cluster assignments and known categorical labels using:

- Adjusted Rand Index (ARI): measures similarity between two labelings, corrected for chance.
- Normalized Mutual Information (NMI): measures the shared information between cluster assignments and known labels.

To further characterize whether clusters were driven by specific annotations (e.g., dominated by single perturbation or cell line), we computed:

- Cluster dominance (≥10% threshold): For each cluster, we calculated the proportion of samples belonging to each label value. We considered a cluster dominant for a label if the most frequent value accounted for at least 10% of cluster members. We reported the fraction of clusters meeting this criterion.
- Cluster purity. For each cluster, we computed the fraction of samples belonging to the most common label value and averaged this quantity across clusters to obtain an overall purity score (range 0-1).

To quantify how strongly each metadata label structured the representation space, we computed an ANOVA-style between-group variance fraction. For a given representation and categorical label, we calculated the total sum of squares relative to the global mean and the between-group sum of squares relative to group means. The ratio of between-group to total variance provided a measure of label-driven structure.

Finally, we computed the Average Silhouette Width (ASW) to evaluate how well samples sharing the same label formed compact and well-separated groups within each representation space.

We performed these analyses across multiple metadata annotations, including perturbation label, replicate, dose, time point, cell line, perturbation type, and MoA.

#### Nearest neighbor search

We computed nearest neighbors using FAISS^57^ with GPU acceleration for efficient similarity search at scale. Prior to indexing, we L2-normalized all vectors so that inner-product similarity corresponded to cosine similarity. We built an exact FAISS index using an inner-product flat index (IndexFlatIP). For each query sample, we retrieved the top 1,000 nearest neighbors. When evaluating retrieval performance, we excluded the first neighbor (self-match).

We evaluated nearest neighbor retrieval using the following metrics for *k ∈ {1,5,10,100}*:

- Hit@k: We considered a query correct if at least one of the top-*k* neighbors shared the same label as the query. We reported Hit@k as the percentage of queries with at least one correct match in the top-*k*.
- Precision@k: For each query, we computed the proportion of retrieved neighbors among the top-*k* that shared the same label as the query and averaged this value across queries. To account for perturbations with limited replicates, we normalized by *min(k,Ri),* where *Ri* denotes the number of available relevant samples for query *i*.

We computed retrieval metrics independently for each representation space and model variant. We evaluated retrieval performance at multiple annotation levels, including perturbation identity (*pert_id*), MoA for compounds, and Gene Ontology (GO) biological process terms for genetic perturbations.

#### Supervised classification of perturbation identity

We trained a multi-class classifier to predict perturbation labels from either Gex profiles or scGPT-derived embeddings. We used the same training/validation split employed during fine-tuning to ensure that validation profiles were never seen during model training.

We implemented a fully connected multilayer perceptron (MLP) in PyTorch. The network consisted of three hidden layers, each comprising a linear transformation followed by ReLU activation and layer normalization. We set the hidden dimensionality equal to the input dimensionality (512 for embeddings; 978 for Gex). The final output layer is projected to *C* classes, corresponding to the number of unique perturbation identifiers (*pert_id*) in the dataset. We trained the model using cross-entropy loss and the Adam optimizer with a batch size of 1,024 for up to 50 epochs. We applied early stopping with a patience of 3 epochs based on validation loss.

After training, we evaluated the classifier on the validation set and reported overall top-1 classification accuracy using *accuracy_score* between predicted and true labels. To assess performance across experimental modalities, we further stratified validation profiles by perturbation type (e.g., compound, shRNA, overexpression) and computed per-category accuracy.

#### Recapitulation of orthogonal information as a classification task (ROC analysis)

We formulated several binary classification tasks based on pairwise similarity. For each task, we defined positive and negative pairs and assessed whether transcriptomic similarity predicted these relationships. Given a pair of compounds (or genes), we used transcriptomic similarity between their representations (defined as 1 - cosine distance) as a scoring function. We computed receiver operating characteristic (ROC) curves and quantified performance using the area under the ROC curve (AUROC). To avoid bias due to unequal numbers of profiles per compound or gene, we performed repeated subsampling. For each of *n=10* iterations (random seed equal to the iteration index), we randomly sampled one profile per compound (or gene).

We evaluated three tasks:

- Chemical similarity: we computed Tanimoto similarities using ECFP4 descriptors (i.e. Morgan fingerprints) for all compounds. We defined chemically similar pairs as those within the top 0.1% of the Tanimoto similarity distributions (i.e., the most chemically similar pairs). We treated all remaining pairs as negatives.
- Target profile similarity (Chemical Checker B4 space): We used Chemical Checker B4 signatures^13,42^, which encode compound-target binding profiles (features correspond to target genes with discrete binding levels). We computed cosine similarities between B4 signatures and defined positives as pairs within the top 0.1% of the similarity distribution. We treated remaining pairs as negatives.
- Genetic perturbation consistency (same gene, different probes): For genetic perturbations (shRNA), we defined positive pairs as profiles derived from different probes targeting the same gene and negative pairs as profiles targeting different genes.

For each task and representation (gene expression profiles and scGPT embeddings), we computed ROC curves and reported AUROC values averaged across the 10 subsampling iterations.

### Knowledge extraction from the perturbational space

#### Prediction of Mechanism-of-Action annotations by nearest-neighbor label propagation

For each unannotated compound, we identified its nearest neighbors with annotated targets using FAISS^57^ with GPU acceleration. We retrieved the top 50 nearest annotated samples based on cosine similarity. To avoid over-representation of compounds with multiple replicate profiles, we collapsed sample-level neighbors to the compound level by retaining only the closest sample per annotated compound. We only generated predictions for query samples with at least three unique annotated compounds among their retained neighbors.

We assigned weights to each annotated compound based on similarity to the query using a softmax transformation of cosine similarities. Using these weights, we computed a probability distribution over MoA labels. For compounds associated with multiple MoA annotations, we distributed their contribution equally across all associated MoAs. We normalized the resulting sample-level MoA probabilities to sum to one. We then aggregated predictions at the compound level by averaging MoA probabilities across all samples belonging to the same unannotated compound. We assigned the final MoA label as the class with the highest mean probability. To quantify consistency across replicate samples, we computed a compound-level agreement score defined as the fraction of samples whose top predicted MoA matched the final compound-level assignment.

To evaluate prediction reliability and calibrate confidence scores, we performed a benchmark using annotated compounds. For each compound belonging to MoA classes with multiple members, we iteratively removed all its profiles from the dataset and predicted its MoA using the same nearest-neighbor procedure.

We then assessed the relationship between confidence score and prediction accuracy, which was used to define a confidence threshold for high-confidence predictions. To restrict predictions to chemically meaningful entities, we applied additional filtering criteria. We retained only compounds with valid SMILES strings, excluded PAINS, and applied physicochemical constraints (150 < molecular weight < 600, 7 < heavy atoms < 55, QED > 0.3). This filtering step resulted in a final set of 22,374 compounds (∼70% of the initially unannotated set).

The final output table included, for each compound: predicted MoA, the confidence score (mean probability of the assigned MoA), agreement between compound samples, and average TAS for each unannotated compound (Supplementary Table 2). We obtained TAS values from the *siginfo_beta.txt* metadata table and aggregated across samples for each compound.

#### Linking chemical and genetic perturbations for target prioritization

For each compound profile, we computed nearest-neighbor shRNA embeddings using FAISS^57^ with GPU acceleration and cosine similarity. We retrieved the top-ranked shRNA neighbors and collapsed them to a ranked list of unique target genes. For each compound-gene pair, we computed a robustness score defined as the fraction of compound profiles for which shRNAs targeting gene *g* appeared among the top 100 nearest shRNA neighbors. This procedure generated, for each compound, a ranked list of candidate target genes ordered by robustness.

We first evaluated a curated benchmark of 13 compound-target pairs from LINCS^6^. For each benchmark compound, we measured the rank and percentile position of the expected target gene in the compound-specific ranked gene list. As a null model, we repeated the full robustness analysis after randomly permuting the mapping between shRNA profiles and target genes. We performed this label-shuffling procedure 200 times. For each permutation, we recomputed target-gene ranks and percentiles, generating a null distribution for comparison with the observed recovery.

We further evaluated target recovery in two larger compound-target benchmark sets. The first was derived from Chemical Checker B4 target annotations^13^, for which we mapped 3,837 compounds and 1,331 targets to the LINCS perturbational space. The second was derived from FRoGS^15^, using Broad compound-target annotations curated by the authors, comprising 2,175 compounds and 1477 targets. For each benchmark, we evaluated whether at least one annotated target appeared among the top 100 highest-ranked genes for each compound.

To identify factors associated with successful target recovery, we performed an exploratory analysis of compound-level profile quality and prediction performance in the Chemical Checker and FRoGS benchmarks. For each compound, we summarized transcriptional activity by aggregating TAS values across all available QC-passing compound profiles and quantified profile support as the number of QC-passing profiles available for that compound. We then assessed whether these variables were associated with target prioritization performance by computing Spearman correlations between each feature and the best rank of any annotated target in the compound-specific ranked gene list. In addition, we classified compounds as recovered or non-recovered according to whether at least one annotated target appeared among the top 100 predicted genes, and we compared TAS and profile-support distributions between these groups.

To prioritize reliable compound-target associations, we computed a compound-level confidence score integrating four features: number of samples, transcriptional activity, functional enrichment coherence, and gene-retrieval robustness. For the number of samples score, we quantified the number of QC-passing compound profiles per compound, log-transformed this value to reduce the influence of highly replicated compounds, and min-max normalized it. For the transcriptional activity score, we obtained TAS values from *siginfo_beta.txt*, aggregated them across profiles for each compound, and min-max normalized the mean TAS. For the functional enrichment coherence score, we tested whether the top-ranked genes for each compound formed coherent biological programs. We used GO Biological Process 2026^65^, KEGG 2026^66^, and Reactome 2024^67^ gene sets loaded from Enrichr GMT files^68^. We excluded terms containing fewer than 10 or more than 500 genes. For each compound-term pair, we computed gene overlap and assessed enrichment using a hypergeometric test. For each compound, we retained the strongest enrichment signal, defined as the lowest raw enrichment p-value across terms, and adjusted these best p-values across compounds using the Benjamini-Hochberg correction. We converted enrichment significance into a normalized score for integration into the composite confidence score. For the gene-retrieval robustness score, we computed the mean robustness of the top 100 compound-gene pairs for each compound. We calculated the final composite confidence score as the arithmetic mean of the four normalized scores.

For Chemical Checker and FRoGS benchmarks, we defined recovered compounds as those with at least one known target among the top 100 predicted genes. We compared score distributions using a one-sided Mann-Whitney U test. We then examined the cumulative fraction of recovered compounds as a function of the composite score.

We finally applied this framework to all compounds in the embedding space.

### Contextualization of new data into the perturbation space

#### Contextualization of unseen conditions compounds and cellular context

We designed two hold-out settings: (i) unseen compounds and (ii) unseen cellular contexts. For the unseen compound setting, we selected 50 compounds spanning diverse Mechanisms-of-Action (MoAs) and excluded all associated gene expression profiles from the training set. These compounds were structurally diverse and representative of the chemical heterogeneity observed in the training set (Supplementary Figure 11). For the unseen cellular context setting, we excluded all perturbation profiles derived from a single cell line (A549) during training. In both cases, we fine-tuned a new HQ model using the same training procedure as in the original setting, for 50 epochs, but without the held-out samples. After training, we generated embeddings for the held-out profiles using *scg.tasks.embed_data()*.

We computed cosine distances between held-out embeddings and embeddings from the training set. For each query profile (either from held-out compounds or from A549), we retrieved the top-k nearest neighbors from the training set using FAISS. We assessed contextualization performance by determining whether at least one of the retrieved neighbors shared the same annotation as the query. For compounds, we evaluated recovery at the MoA level, whereas for the cell-line hold-out setting, we evaluated recovery both at the exact perturbation level and at the MoA level. We also computed the probability of recovering at least one label-consistent neighbor under random sampling. For each query *i*, we defined *N* as the total number of reference profiles, *X_i_* as the number of reference profiles sharing the same annotation as the query, and *k* as the number of retrieved neighbors. We computed the probability of obtaining at least one matching neighbor under sampling without replacement and averaged this probability across all queries to estimate the expected hit rate.

We performed K-means clustering on the training embedding space. We set the number of clusters as the square root of the number of training samples. We assigned held-out embeddings to their nearest cluster centroid. For each cluster, we tested enrichment of label annotations using a hypergeometric test with Benjamini-Hochberg correction. We considered a held-out profile correctly contextualized if it was assigned to a cluster significantly enriched (false discovery rate < 0.05) for at least one of its true annotations.

To evaluate whether limited exposure to a new cellular context improves contextualization, we tested two adaptation strategies. First, we fine-tuned a model excluding A549 perturbation profiles while retaining A549 control (DMSO) profiles. Second, we adapt the fine-tuning model by training 15 more epochs using a reconstruction objective on A549 control profiles. We compared both strategies against the non-adapted model.

#### Integrate new data into embedding space

We selected an external gene expression dataset from GEO (GSE51212)^60^, which contains transcriptional profiles of HCC827 lung cancer cells treated with three compounds: erlotinib, AZD-6244, and BEZ-235. Each compound was profiled in biological triplicates at a concentration of 1 µM after 6 hours of treatment. In addition, erlotinib was measured at multiple time points (3, 12, and 24 hours). All three compounds are also present in the LINCS dataset, enabling direct comparison between external and reference perturbation profiles.

We generated embeddings for all external gene expression profiles using the high-quality fine-tuned scGPT model, following the same preprocessing and embedding extraction procedure applied to LINCS data. We then computed cosine distances between external embeddings and all embeddings in the LINCS perturbational space. For each external profile, we ranked LINCS embeddings according to similarity and recorded the rank position at which the corresponding compound (same perturbagen identity) was recovered.

## Data availability

All datasets analyzed in this study are publicly available. LINCS L1000 Level 3 data^6^ (2020 release) can be obtained from the CLUE data library (https://clue.io/data). The external dataset used for contextualization is available from the Gene Expression Omnibus (GEO) under accession GSE51212^60^. Chemical Checker^13^ compound-target annotations and the gene set collections used in the analyses are provided through our GitHub repository (https://github.com/sbnb-irb/LINCS_scGPT_embeddings), whereas FRoGS^15^ compound-target annotations are available from the original repository (https://github.com/chenhcs/FRoGS/blob/main/data/cpd_gene_pairs.cs). The fine-tuned scGPT models and all LINCS perturbation embeddings generated in this study are available through Zenodo (https://zenodo.org/records/20925618).

## Code availability

All code developed for this study is publicly available at https://github.com/sbnb-irb/LINCS_scGPT_embeddings. The repository includes scripts and Jupyter notebooks for LINCS preprocessing, scGPT fine-tuning, embedding generation, nearest-neighbor analyses, mechanism-of-action inference, compound-target prioritization, and reproduction of all figures and results presented in the manuscript. Detailed documentation describing the workflow, required input data, and repository structure is also provided.

## Acknowledgments

We would like to thank all public databases and methods that have allowed this study, and the members of the Structural Bioinformatics and Network Biology lab from IRB Barcelona for the constant discussions.

## Funding

PA acknowledges the support of the Generalitat de Catalunya (2021 SGR 00876), the Spanish Ministerio de Ciencia, Innovación y Universidades (PID2023-152296OB-I00), and the European Commission (CLARITY: 101137201). EP-L has been a recipient of an FI fellowship (2022 FI_B_00767), which is co-financed by the European Union through the European Social Fund Plus (ESF +). We also acknowledge institutional funding from the Spanish Ministry of Science and Innovation through the Centres of Excellence Severo Ochoa Award, and from the CERCA Programme / Generalitat de Catalunya.

## Author information

### Authors and Affiliations

Institute for Research in Biomedicine (IRB Barcelona), The Barcelona Institute of Science and Technology (BIST), Barcelona, Catalonia, Spain

### All the authors

Institució Catalana de Recerca i Estudis Avançats (ICREA), Barcelona, Catalonia, Spain Patrick Aloy

## Contributions

EP-L and PA conceived and designed the study, analyzed the results and wrote the manuscript. EP-L implemented all the computational models. All authors read and approved the final manuscript.

### Corresponding author

Correspondence should be addressed to Patrick Aloy.

## Competing interests

The authors declare no competing interests.

## Source data

All the experimental data generated in this study and the source data to reproduce the analysis are publicly available. See *Data Availability*.

## Supplementary Figures

**Supplementary Figure 1.**
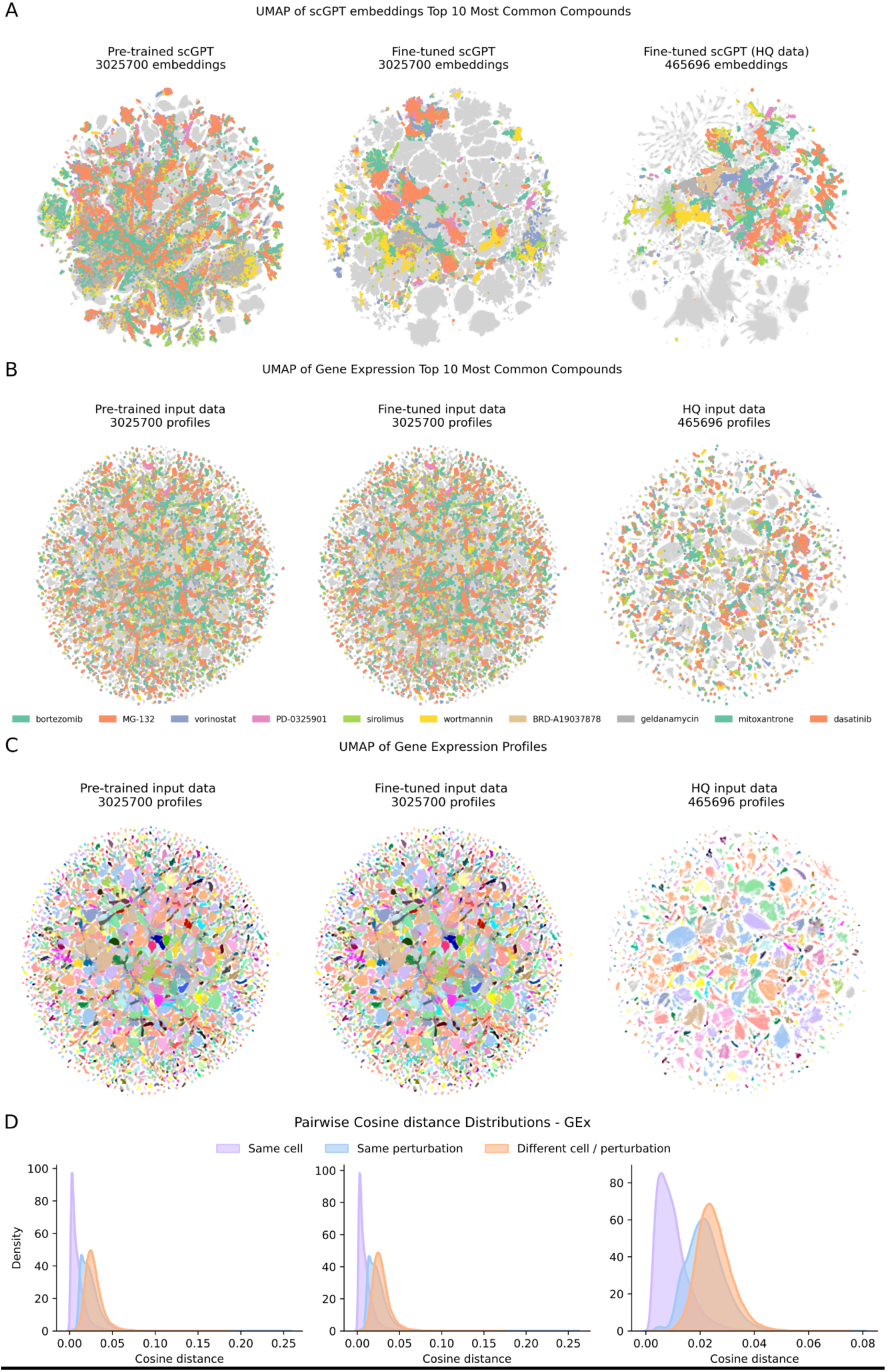
Exploration of perturbation structure in gene expression and embedding spaces. **A, B.** UMAP representations of the scGPT embeddings (A) and gene expression (Gex) profiles (B) derived from the LINCS L1000 Level 3 dataset. Colors indicate the 10 most abundant perturbations. **C.** UMAP representation of LINCS gene expression profiles colored by cell line of origin. **D.** Pairwise cosine distance distributions for gene expression profiles in the three datasets used to generate the embeddings (pre-trained, full, and HQ), grouped by same cell line (purple), same perturbation (blue), or random pairs (orange).

**Supplementary Figure 2.**
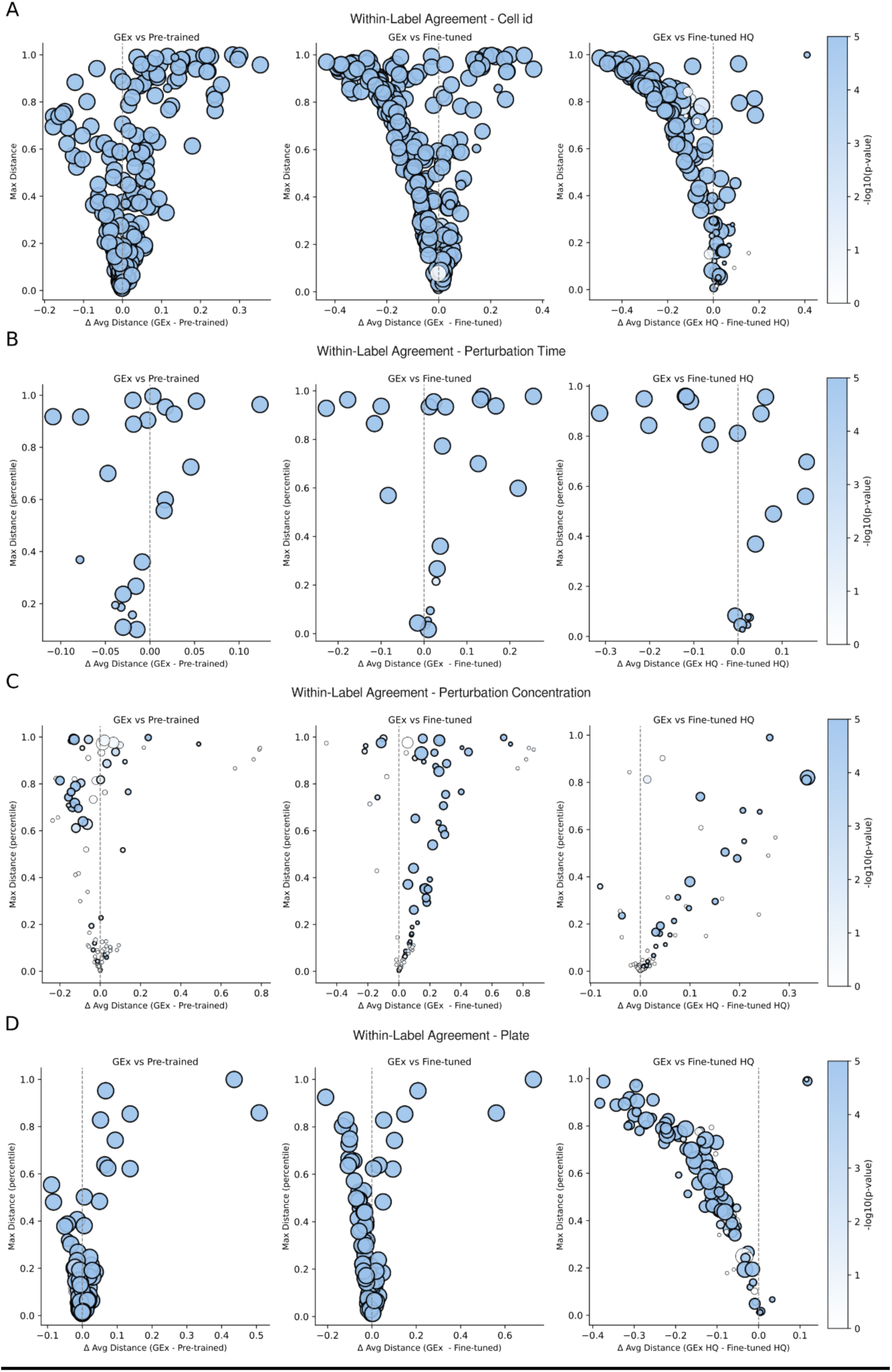
Intra-coherence analysis across different metadata features. Each point represents a cell line **(A)**, perturbation time point **(B)**, concentration **(C)**, or plate **(D)**. The x-axis shows the difference in average percentile-normalized within-category distance between gene expression (Gex) profiles and scGPT embeddings (left = tighter in Gex; right = tighter in scGPT embeddings), while the y-axis shows the maximal within-category distance. Point size reflects the number of samples within each category, and point color (-log10 *p*-value) indicates the significance of the difference between representations.

**Supplementary Figure 3.**
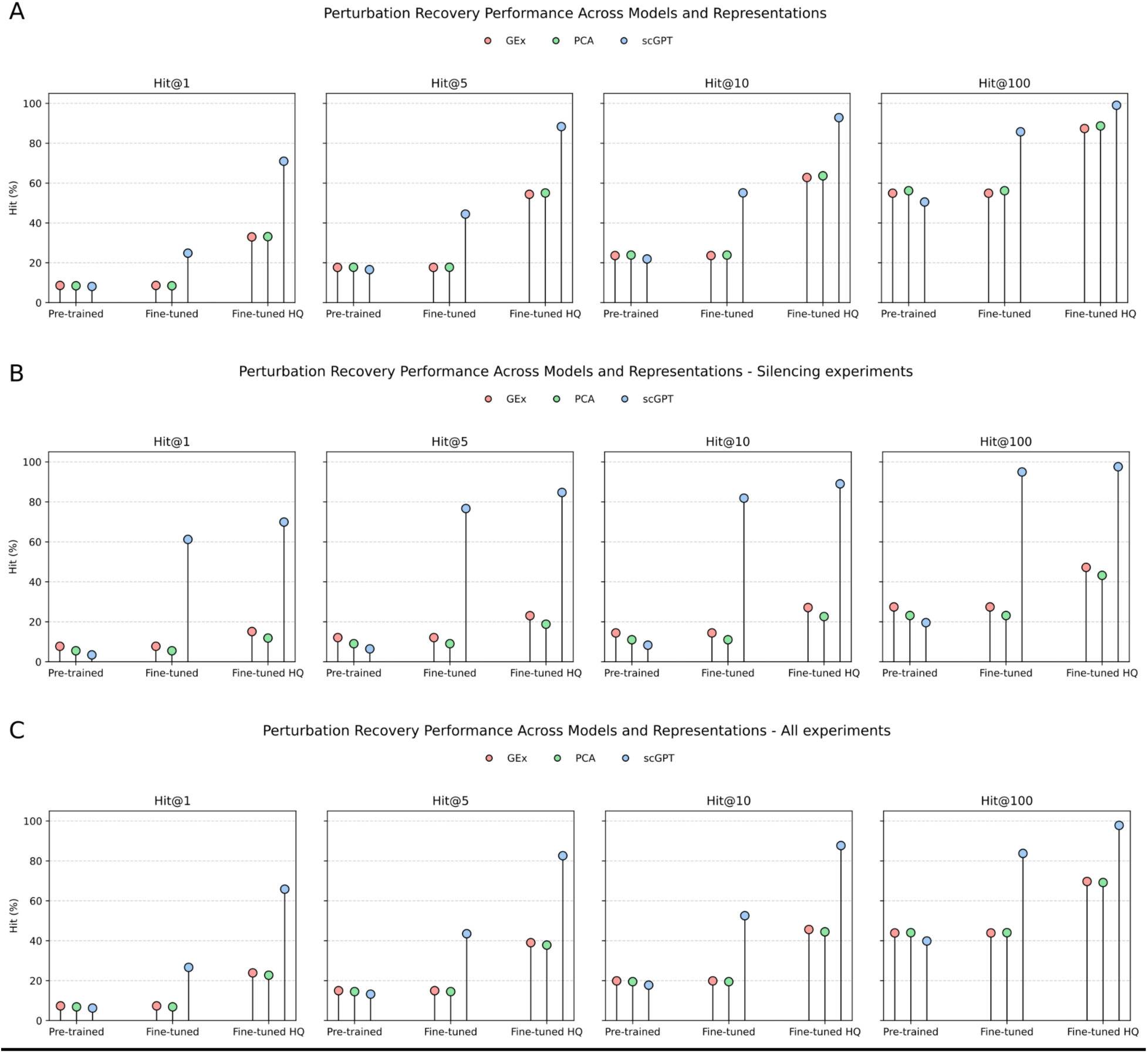
Perturbation recovery analysis. Percentage of perturbations for which at least one nearest neighbor shared the same perturbation label among the top 1, 5, 10, or 100 nearest neighbors (Hit@k) in gene expression (Gex) (red), PCA (green), or scGPT embedding (blue) spaces across the pretrained, full fine-tuned, and high-quality (HQ) fine-tuned models. Results are shown for compound perturbations **(A)**, gene silencing perturbations **(B)**, and all perturbations combined **(C)**.

**Supplementary Figure 4.**
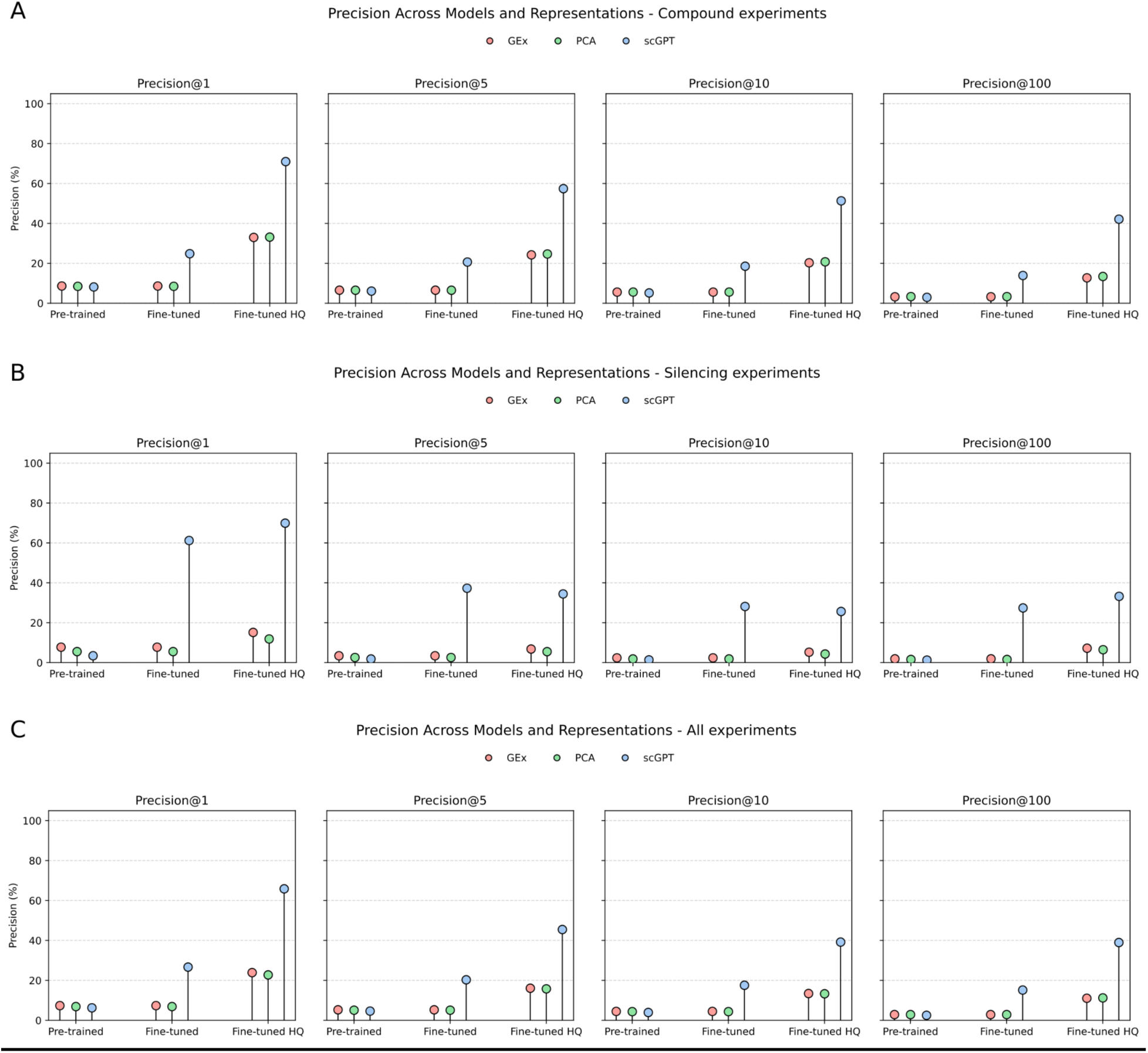
Perturbation precision analysis. Precision@k for perturbation label recovery in gene expression (Gex) (red), PCA (green), and scGPT embedding (blue) spaces across the pretrained, full fine-tuned, and high-quality (HQ) fine-tuned models. Precision was computed for the top 1, 5, 10, and 100 nearest neighbors as the proportion of retrieved neighbors sharing the same perturbation label as the query. Results are shown for compound perturbations **(A)**, gene silencing perturbations **(B)**, and all perturbations combined **(C)**.

**Supplementary Figure 5.**
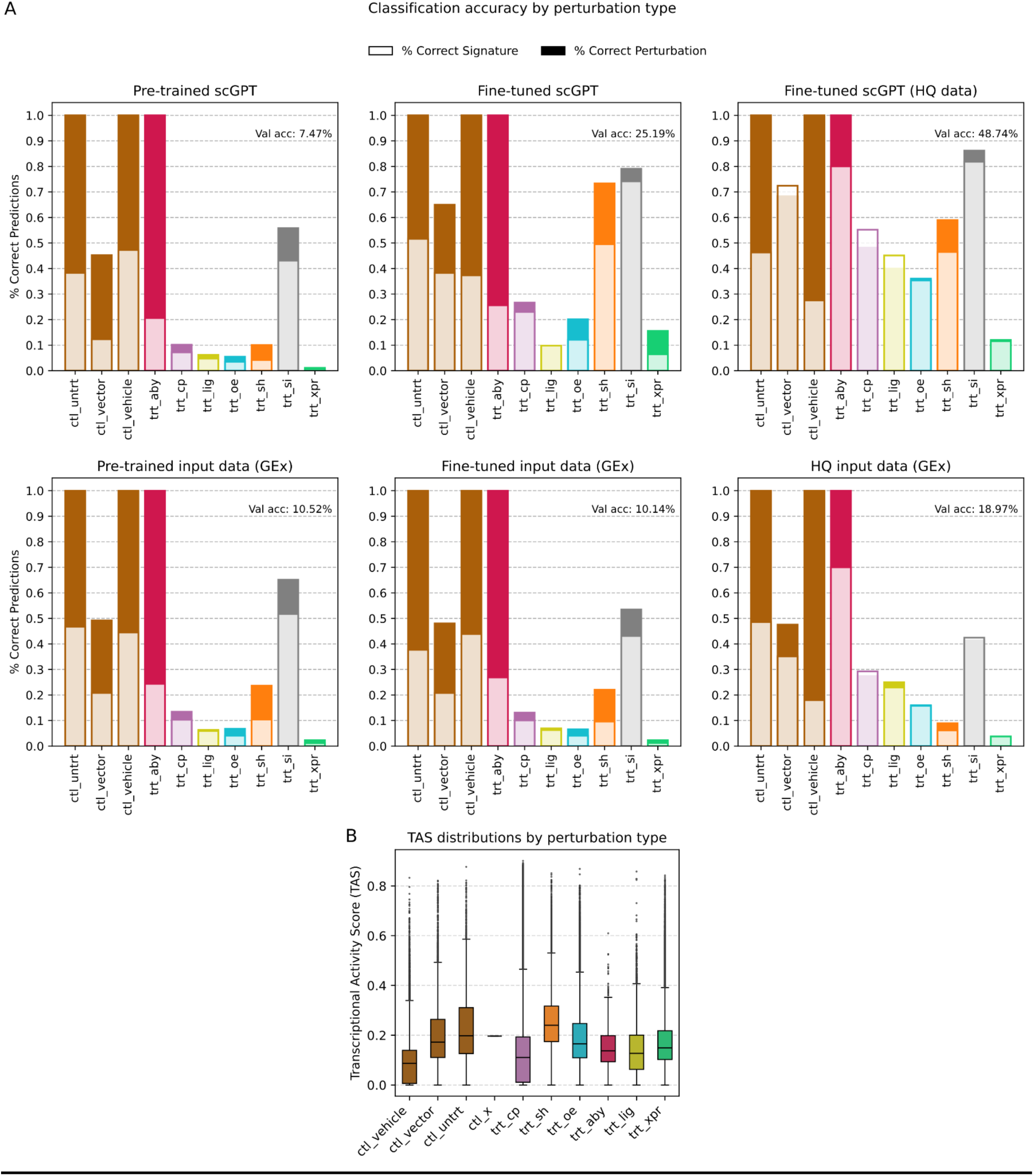
Classification performance stratified by perturbation type. **A.** Classification accuracy by perturbation type for perturbation identity prediction. Top panels show results obtained using scGPT embeddings, whereas bottom panels show results obtained using gene expression (Gex) profiles. Dark bars represent the percentage of correctly classified profiles, and light bars represent the percentage of correctly classified unique perturbations. **B**. Distribution of Transcriptional Activity Scores (TAS) across perturbation types.

**Supplementary Figure 6.**
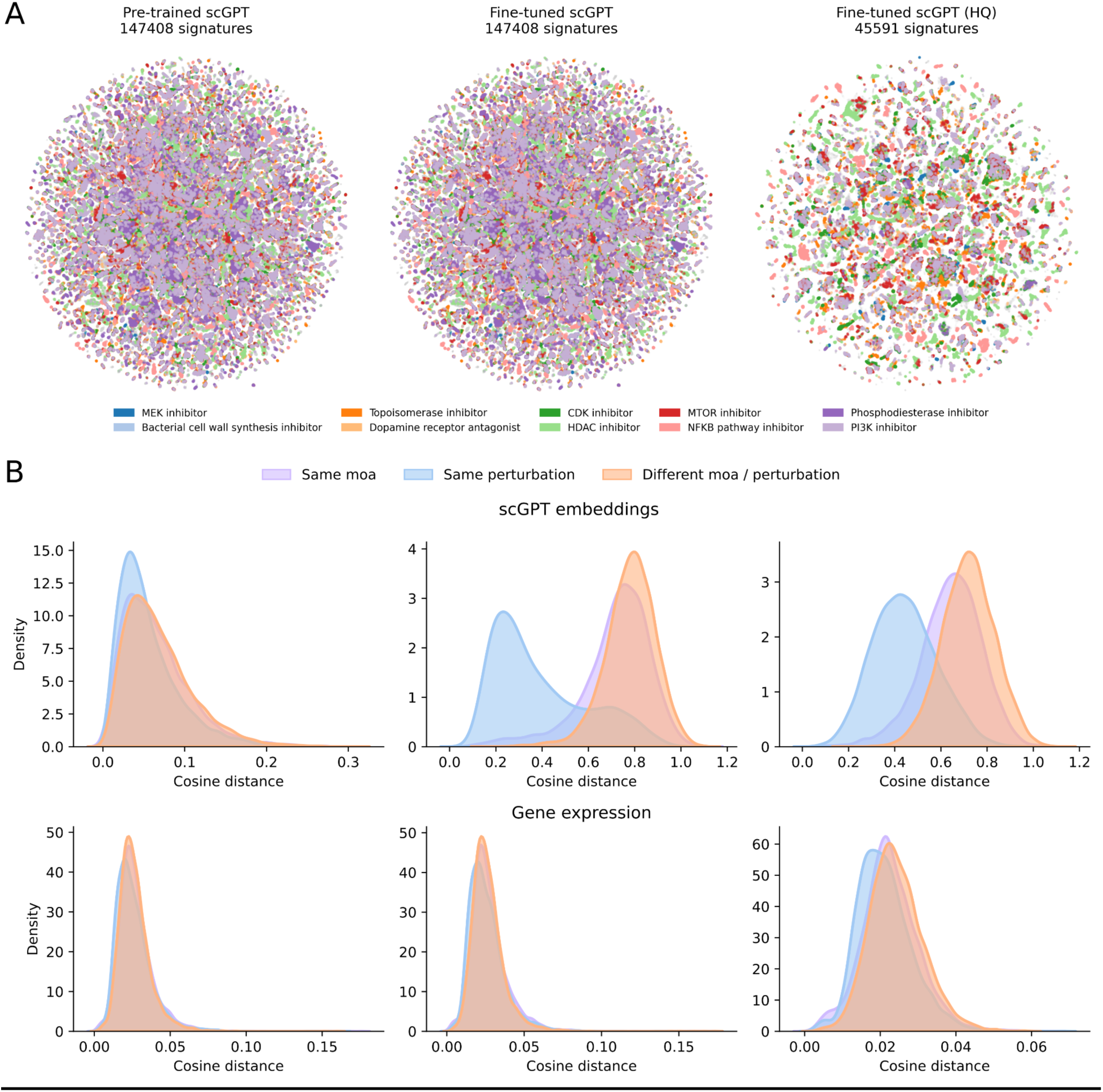
Exploration of orthogonal biological information in gene expression and embedding spaces. **A.** UMAP representation of gene expression (Gex) profiles colored by the most abundant Mechanism-of-Action (MoA) classes. **B.** Pairwise cosine distance distributions for scGPT embeddings (top row; pre-trained, full fine-tuned, and HQ fine-tuned spaces) and gene expression profiles (bottom row; pre-trained, full, HQ), grouped by same perturbation (blue), same MoA (purple), or random pairs (orange)

**Supplementary Figure 7.**
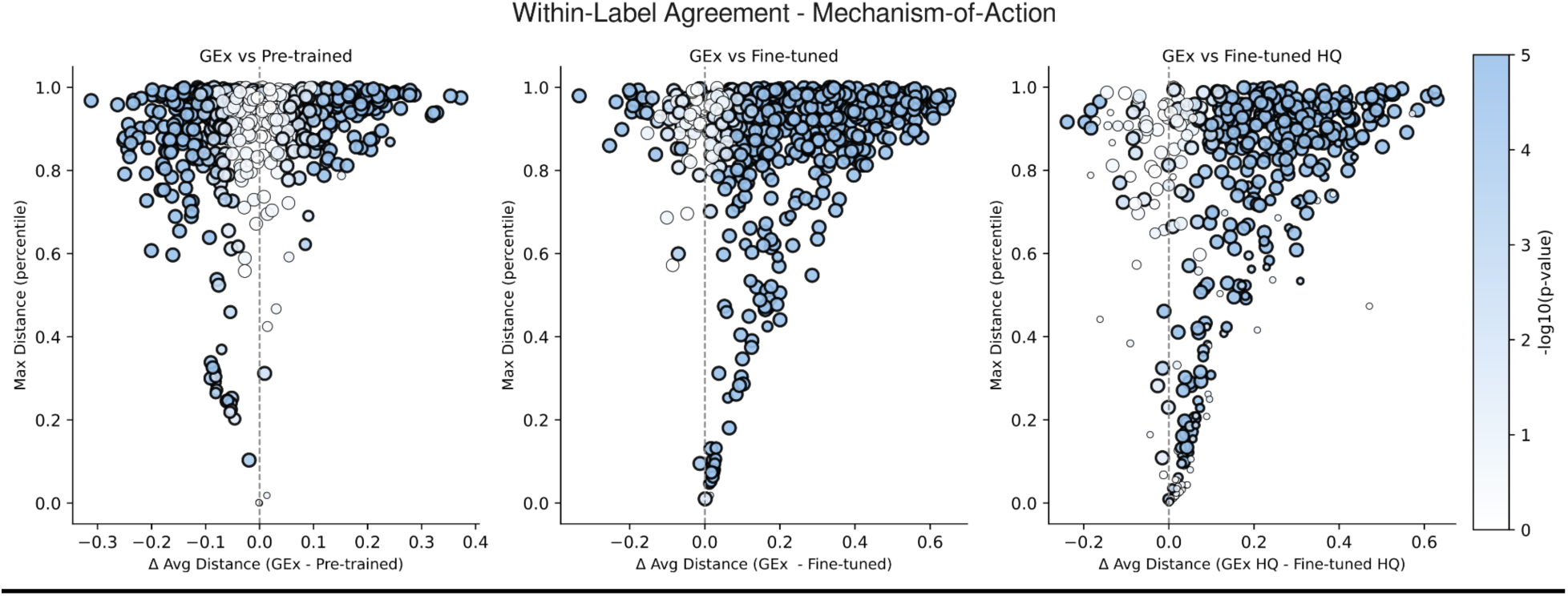
Intra-coherence analysis for Mechanism-of-Action (MoA) classes. Each point represents a MoA class. The x-axis shows the difference in average percentile-normalized within-MoA distance between gene expression (Gex) profiles and scGPT embeddings (left = tighter in GEx; right = tighter in scGPT embeddings), while the y-axis shows the maximal within-MoA distance. Point size reflects the number of samples within each MoA class, and point color (-log10 *p*-value) indicates the significance of the difference between representations.

**Supplementary Figure 8.**
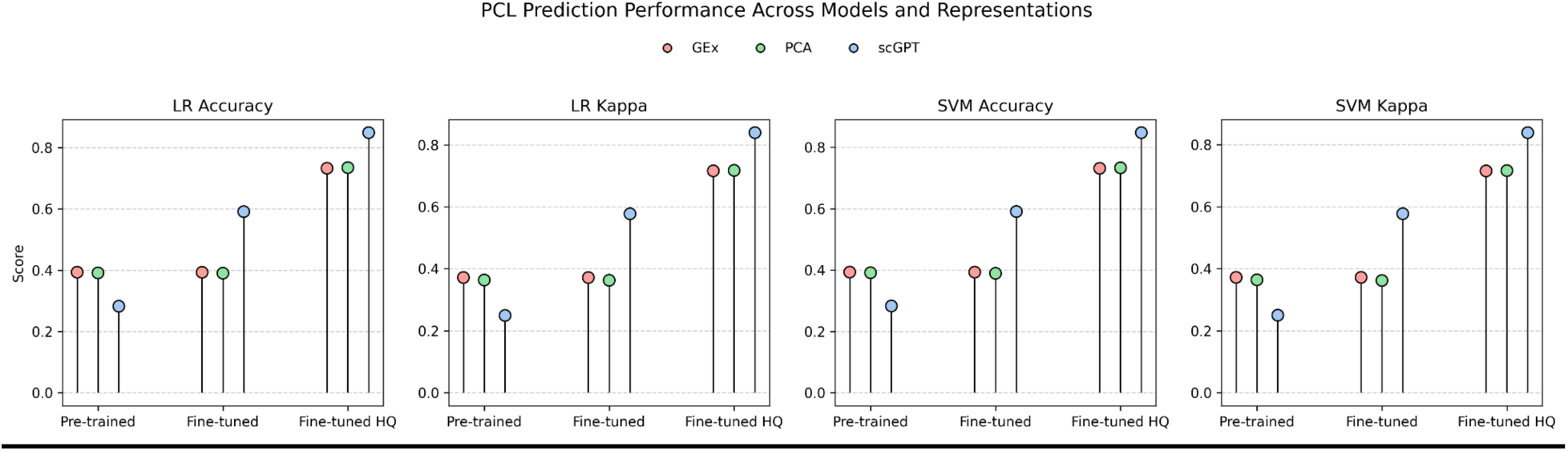
Prediction performance for perturbagen classes (PCLs). We evaluated perturbagen class prediction using logistic regression (LR) and support vector machine (SVM) classifiers across the three model variants (pre-trained, full fine-tuned, and high-quality (HQ) fine-tuned) and the three data representations (gene expression (Gex) (red), PCA (green), and scGPT embeddings (blue)). We report classification accuracy and Cohen’s kappa score.

**Supplementary Figure 9.**
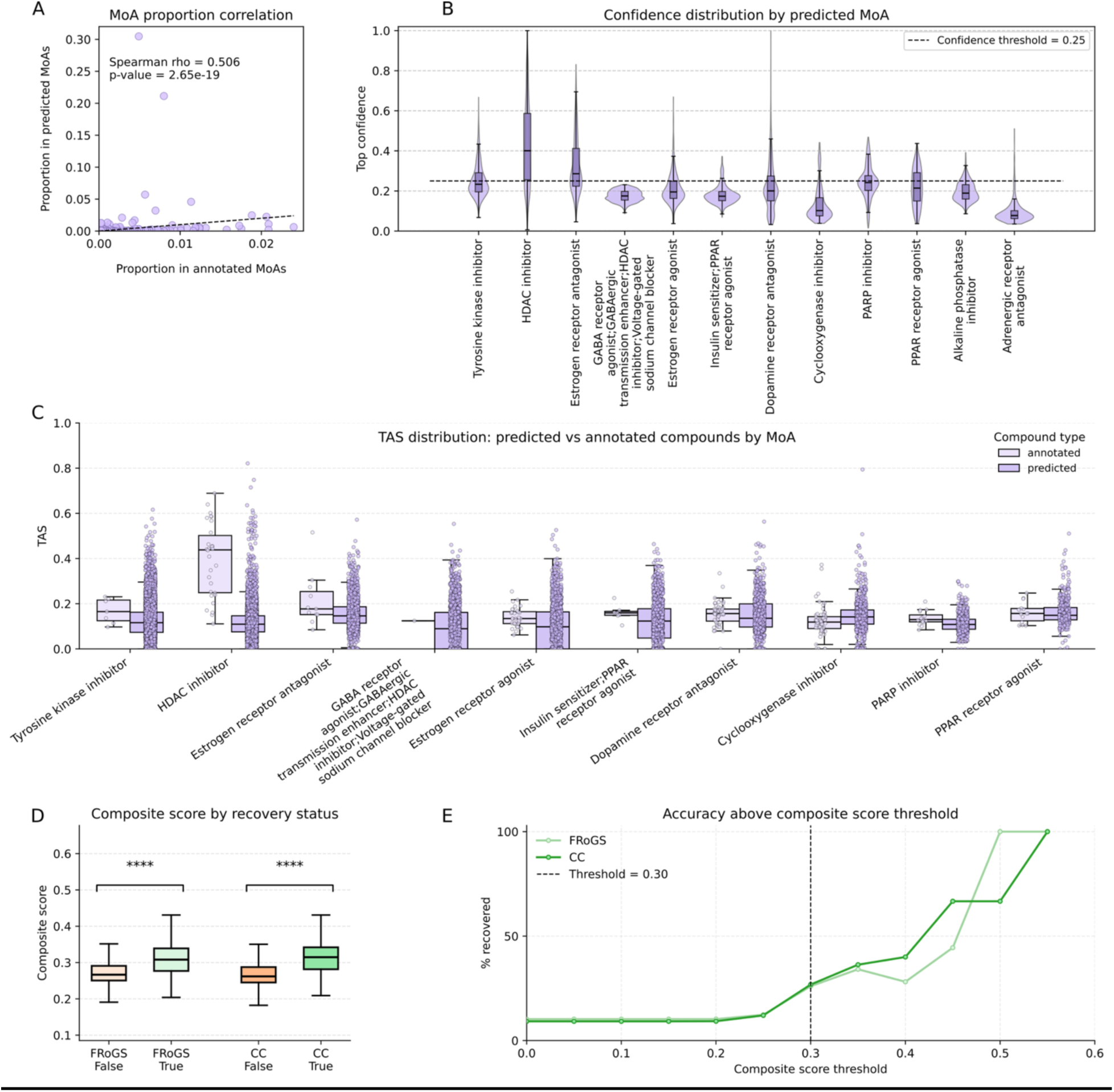
Evaluation and calibration of confidence scores for MoA and target prioritization analyses. **A.** Correlation between the proportion of annotated samples assigned to each Mechanism-of-Action (MoA) class and the proportion of unannotated samples predicted to belong to the same MoA class. **B.** Distribution of confidence scores across predictions for the 12 most frequently predicted MoA classes. The selected confidence threshold (0.25) is indicated by the dashed line. **C.** Distribution of compound-level TAS for annotated and predicted compounds across the top predicted MoA categories. **D.** Distribution of composite confidence scores stratified by target recovery status (recover green; not recover orange) across the Chemical Checker and FRoGS benchmark datasets. **E.** Calibration of the composite confidence score for target prioritization. Curves show the cumulative proportion of compounds for which at least one correct target was recovered within the top 100 predicted genes in the FRoGS (light green) and Chemical Checker (dark green) benchmark datasets as a function of the composite confidence score. The selected confidence threshold (0.35) is highlighted.

**Supplementary Figure 10.**
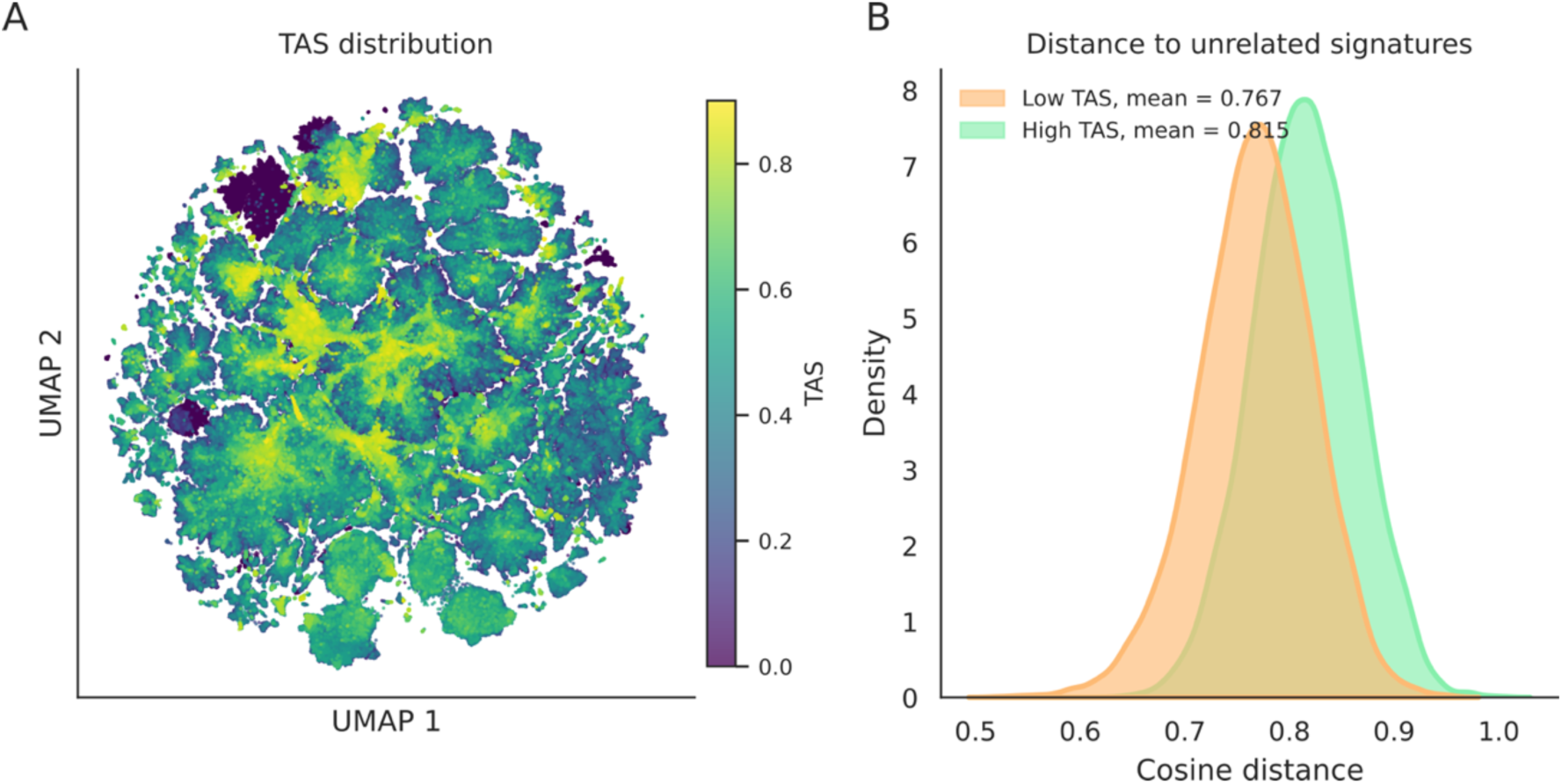
Spatial proximity of low-TAS embeddings. **A.** UMAP visualization of scGPT embeddings colored by transcriptional activity score (TAS). Points are plotted in ascending TAS order, so high-TAS embeddings are shown on top. **B.** Average cosine distance from low- and high-TAS embeddings to random unrelated embeddings. Low-TAS and high-TAS groups correspond to the bottom 5% and top 5% of TAS values, respectively. For each group, 10,000 embeddings were randomly selected. For each selected embedding, up to five random unrelated embeddings were sampled, excluding embeddings from the same compound, or any shared MoA. Cosine distances were averaged per query embedding before comparing the two TAS groups.

**Supplementary Figure 11.**
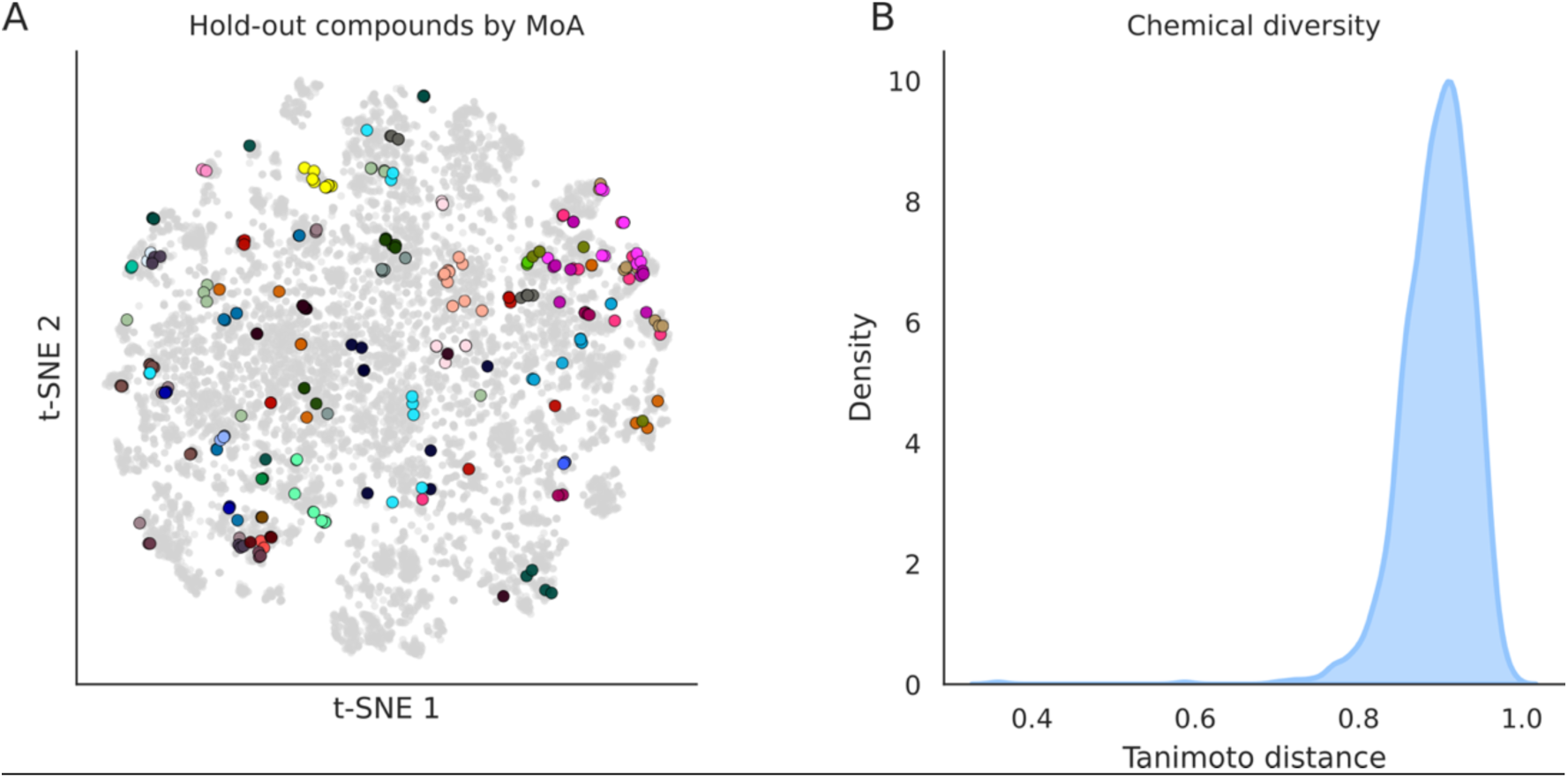
Characterization of held-out compounds and TAS-dependent embedding structure. **A.** t-SNE visualization of scGPT embeddings in the high-quality (HQ) embedding space for compounds with known mechanisms of action (MoAs). A maximum of 10 embeddings per compound was randomly selected. Embeddings corresponding to the held-out compounds are highlighted and colored by MoA. The held-out compound set comprises 51 distinct MoAs. **B.** Pairwise Tanimoto distance between held-out compounds, showing the chemical diversity of the held-out set.

**Supplementary Figure 12.**
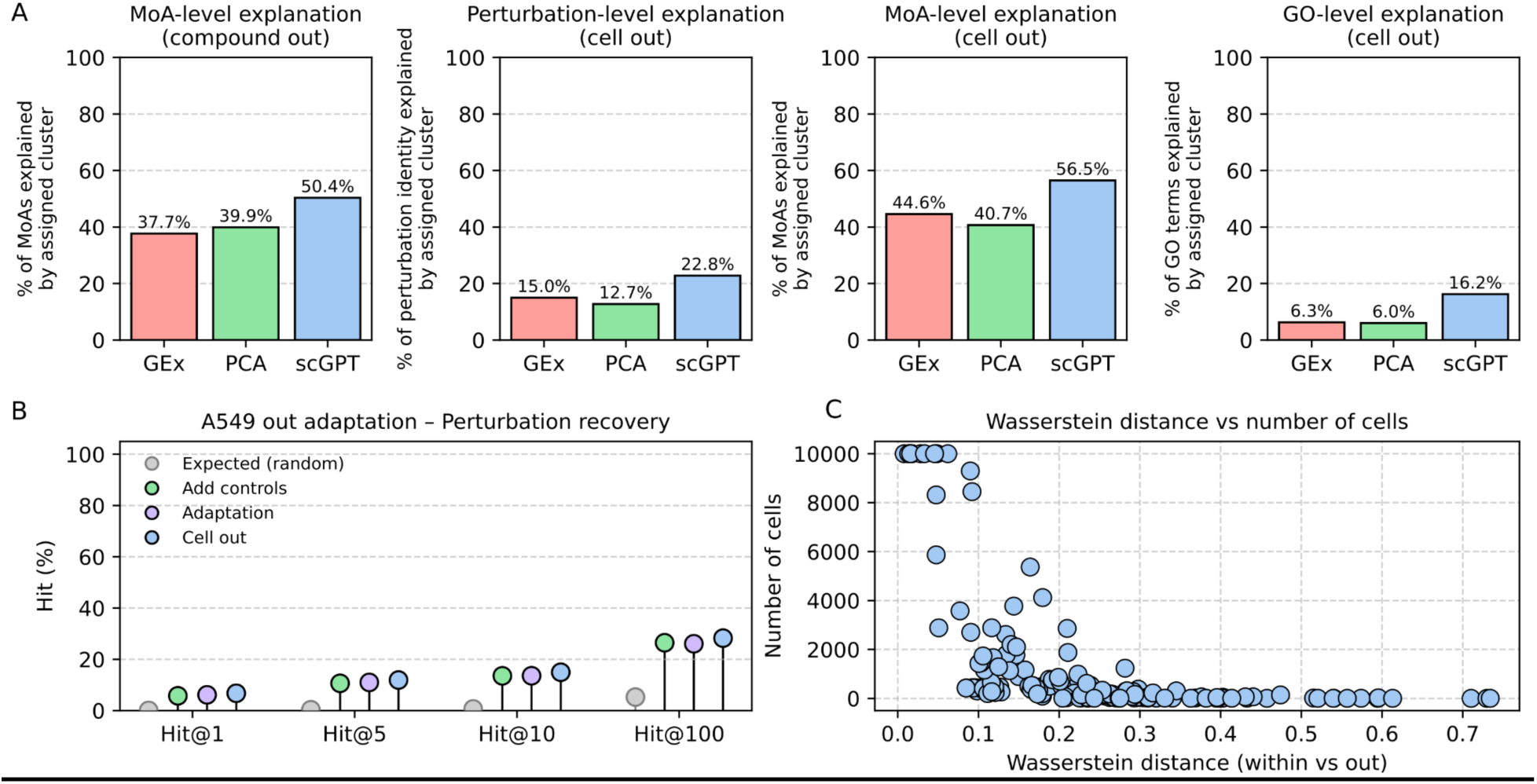
Generalization and contextualization performance across unseen compounds and cellular contexts. **A.** Contextualization performance of HQ fine-tuned scGPT embeddings under two hold-out settings: unseen compounds (left) and unseen cell line (right panels). Bars represent the percentage of held-out profiles correctly assigned to clusters significantly enriched for the corresponding Mechanism-of-Action (MoA), perturbation label, or Gene Ontology (GO) annotation. We compare the performance of using gene expression (Gex) profiles (red), PCA (green), scGPT embeddings (red). **B.** Contextualization performance under the A549 cell line hold-out setting, quantified as Hit@k (k = 1, 5, 10, 100) for exact perturbation label recovery. We compared two adaptation strategies: complete A549 exclusion during training (blue), retention of A549 control profiles only (green), and additional adaptation using a reconstruction loss on A549 controls for 15 epochs (purple). Random expectation is shown in grey. **C.** Wasserstein distance between within-cell-line and between-cell-line distance distributions in the HQ fine-tuned embedding space as a function of the number of profiles available for each cell line. For cell lines with more than 10,000 profiles, we randomly subsampled 10,000 profiles before analysis.

## Notes

### Competing Interest Statement

The authors have declared no competing interest.

